# Cell type- and chromosome-specific chromatin landscapes and DNA replication programs of *Drosophila* testis tumor stem cell-like cells

**DOI:** 10.1101/2025.04.19.649648

**Authors:** Jennifer A. Urban, Daniel Ringwalt, John M. Urban, Wingel Xue, Ryan Gleason, Keji Zhao, Xin Chen

## Abstract

Stem cells have the unique ability to self-renew and differentiate into specialized cell types. Epigenetic mechanisms, including histones and their post-translational modifications, play a crucial role in regulating programs integral to a cell’s identity, like gene expression and DNA replication. However, the transcriptional, chromatin, and replication timing profiles of adult stem cells *in vivo* remain poorly understood. Containing germline stem cells (GSCs) and somatic cyst stem cells (CySCs), the *Drosophila* testis provides an excellent *in vivo* model for studying adult stem cells. However, the small number of stem cells and cellular heterogeneity of this tissue have limited comprehensive genomic studies.

In this study, we developed cell type-specific genomic techniques to analyze the transcriptome, histone modification patterns, and replication timing of GSC-like and CySC-like cells. Single cell RNA sequencing validated previous findings on GSC-CySC intercellular communication and revealed high expression of chromatin regulators in GSC-like cells. To characterize chromatin landscapes, we developed a cell-type-specific chromatin profiling assay to map H3K4me3-, H3K27me3-, and H3K9me3- enriched regions, corresponding to euchromatic, facultative heterochromatic, and constitutive heterochromatic domains, respectively. Finally, we determined cell type-specific replication timing profiles, integrating our *in vivo* datasets with published data using cultured cell lines. Our results reveal that GSC-like cells display a distinct replication program compared to somatic lineages, that aligns with chromatin state differences. Collectively, our integrated transcriptomic, chromatin, and replication datasets provide a comprehensive framework for understanding genome regulation differences between these *in vivo* stem cell populations, demonstrating the power of multi-omics in uncovering cell type-specific regulatory features.

## INTRODUCTION

Stem cells are defined by their ability to self-renew and differentiate into specialized cell types. During development and tissue homeostasis, asymmetric cell division of stem cells produces two genetically identical daughter cells that adopt distinct cell fates. It is hypothesized that asymmetric inheritance of cell fate determinants plays a critical role in specifying these identities. Epigenetic mechanisms, such as histones and their post-translational modifications, regulate chromatin compaction, transcription, and DNA replication timing, to influence cell fate decisions. In stem cells, epigenetic regulation is thought to be a key contributor to their differentiation potential. However, its precise roles in *in vivo* stem cell systems remain to be fully elucidated.

The *Drosophila* male germline is a well-established model for studying stem cell biology with well-defined cell lineages and differentiation programs. Germline stem cells (GSCs) reside near the stem cell niche and undergo asymmetric cell divisions to generate a self-renewing GSC and a differentiating gonialblast (GB, Fig. 1A) (Fuller and Spradling 2007; Matunis et al. 2012). Concurrently, cyst stem cells (CySCs) divide to produce a CySC and a cyst cell (CC) (Cheng et al. 2011; Gönczy and Dinardo 1996). The GB, encapsulated by two CCs, undergoes four mitotic divisions with incomplete cytokinesis, forming a 16-cell spermatogonial (SG) cyst that subsequently enters meiosis (Fig. 1A) (Matunis et al. 2012).

**Fig. 1.**
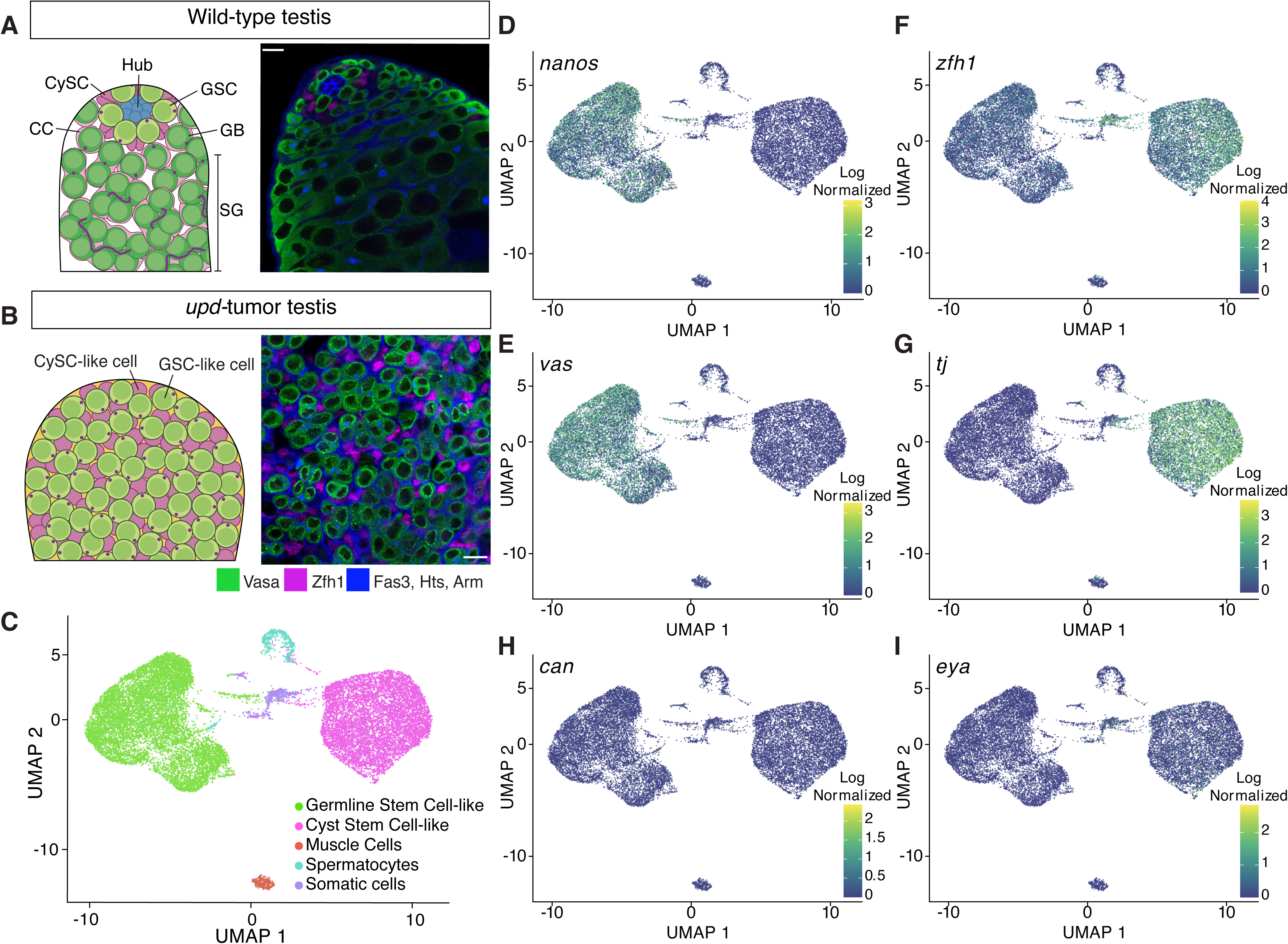

Traditional genetic and cell biological approaches have provided key insights into lineage-specific differences between GSCs and CySCs. These lineages communicate through cell-cell adhesion molecules and signaling pathways to coordinate germline cyst differentiation to become mature sperm (Matunis et al. 2012). However, the cell heterogeneity and small number of stem cells in wild-type testis tissue have made it difficult to generate comprehensive cell-type-specific genomic datasets defining their global transcriptional, chromatin, and replication landscapes. To address this, we developed cell-type-specific genomic strategies for testicular stem cells derived from a tumor model, enabling a detailed analysis of their transcriptome, chromatin landscapes, and replication timing dynamics. This approach uncovered previously unrecognized chromosome-specific differences in chromatin regulation between GSCs and CySCs, shedding new light on the molecular mechanisms regulating stem cell identity and activity.

## RESULTS

### Using a GSC and CySC tumor model for genomic studies

Our understanding of the transcriptional, chromatin, and replication landscapes of *Drosophila* GSCs is limited due to their small number in wild-type testes (Fig. 1A, Supplemental Fig. S1A). GSCs were underrepresented in previous bulk genomic analyses of wild-type or early-stage germ cell-enriched testes (Gan et al. 2010; Vedelek et al. 2018). Furthermore, even single-cell (scRNA-seq) or single-nucleus (snRNA-seq) transcriptomic methods using whole wild-type testes yielded sparse GSC data (Witt et al. 2019; Yu et al. 2023; Raz et al. 2023; Shi et al. 2020).

To overcome this limitation, we used testes enriched with GSC-like and somatic CySC-like cells by over-expressing the Janus kinase (JAK)-signal transducer and activator of transcription (STAT) pathway ligand, Unpaired (Upd) (Fig.1B, Supplemental Fig. S1B). Hyper-activation of the JAK-STAT pathway in the testis results in over-proliferation of both stem cell types (Kiger et al. 2001; Tulina and Matunis 2001; Leatherman and Dinardo 2010), resulting in a more homogenous tissue composition for genomic profiling of stem cells from both germline and somatic lineages.

In our experimental design, we used two different cell type-specific drivers to overexpress Upd in the fly testes. Driving expression of Upd with either a germline (*nanos* or *nanos*-Gal4) or somatic driver (*traffic jam* or *tj*-Gal4) leads to testis tumors with over-proliferative GSC-like and CySC-like cells. A GFP-tagged histone H3 (H3-GFP) was driven along with Upd in either GSC-like cells (*nanos*>*upd*, *H3- GFP*) or CySC-like cells (*tj*>*upd*, *H3-GFP*). This tumor model reduces cellular heterogeneity, making it an ideal system for studying GSC- or CySC-specific transcriptional, chromatin, and replication landscapes.

### scRNA-seq profiling of GSC-like and CySC-like cells

We performed single-cell RNA sequencing (scRNA-seq) on *nanos*- and *tj*-driven Upd- overexpressing tumor samples (*upd*-tumors), with two replicates each. Across these four samples, we consistently identified distinct germline and somatic lineages, demonstrated through single-cell integration, principal component analysis (PCA), and clustering (Supplemental Figs. S1C-E).

After individually clustering each sample, we integrated them to reduce batch effect variance, and subset the data to GSC-like and CySC-like clusters. Consequently, most cells are clustered into two groups: GSC-like cells (expressing *vasa* and *nanos*) or CySC-like cells (expressing *tj* and *zfh1*) (Figs. 1C- G, Supplemental Table 1). Beyond these two main clusters, we identified three additional clusters: primary spermatocytes (*mst87f*-expressing), muscle cells (*act57B-*expressing), and an unidentified *wb-* expressing somatic lineage (Fig. 1C, Supplemental Fig. S1H, Supplemental Table 1) (Kuhn et al. 1991; Kelly et al. 2002). For this study, we focus on the GSC-like and CySC-like clusters.

The GSC-like and CySC-like clusters followed expected gene expression patterns for known cell type- and stage-specific markers. For example, GSC-like cells, but not CySC-like cells, express *nanos* and *vasa* (Figs. 1D-E, Supplemental Table 1). In contrast, CySC-like cells, but not GSC-like cells, express *zfh1* and *tj* (Figs. 1F-G, Supplemental Table 1).

The expression of stage-specific transcripts further confirms that the *upd*-tumor is enriched with stem cell-like germline and cyst cells. As expected, *stat92E* is detected in both clusters, indicating JAK-STAT pathway activation due to *upd* overexpression (Supplemental Fig. S1I, Supplemental Table 1) (Amoyel and Bach 2012). Additionally, as expected for GSCs, neither *bag-of-marbles* (*bam*), which is a marker of spermatogonia, nor *cannonball* (*can*), a spermatocyte-specific transcript, are strongly expressed in the GSC-like cell cluster (Fig. 1H, Supplemental Fig. S1I, Supplemental Table 1) (McKearin and Spradling 1990; Hiller et al. 2001). Consistently, *eyes-absent* (*eya*), a late-stage cyst cell marker, is not expressed in the CySC-like cell cluster (Fig. 1I, Supplemental Table 1) (Fabrizio et al. 2003).

We next compared our scRNA-seq data to published snRNA-seq from whole wild-type testes to investigate whether our results are consistent with the cell clusters identified in the published dataset (Li et al. 2022; Raz et al. 2023). Our analysis confirms that GSC-like cells from the *upd*-tumor are transcriptionally comparable to the wild-type cluster containing GSCs and early spermatogonia (Supplemental Fig. S2). Similarly, CySC-like cells show transcriptional similarity to the annotated cyst stem cell cluster in the published dataset (Supplemental Fig. S2). These comparisons support our conclusion that the two major clusters from the *upd*-tumor correspond to GSCs and CySCs, expressing characteristic cell type- and stage-specific transcripts. The size and purity of these clusters provide an opportunity to robustly profile the transcriptomes of GSC- and CySC-like cells.

### CySC-like cells highly express intercellular communication genes while GSC-like cells enrich for chromatin regulators

To identify cell-type-enriched transcripts, we performed differential gene expression analysis. A transcript was considered as having significantly higher expression in GSC-like cells if it had a fold change >1.5 (GSC/CySC) and an S-value < 10^-4^ (estimated false sign rate) (Fig. 2A, Supplemental Table 2). We identified 827 genes enriched in GSC-like cells and 1,794 in CySC-like cells. These GSC-like and CySC-like enriched gene sets contained expected markers, supporting cluster identity and the robustness of our analysis (Fig. 2B, Supplemental Table 2). First, the long-noncoding RNAs, *roX1* and *roX2* (*RNA on the X*), components of the male dosage compensation complex found in somatic but not germ cells (Shevelyov et al. 2022), rank among the top 100 genes with significantly higher expression in CySC-like cells (Fig. 2B, Supplemental Table 2). Second, *ago3*, encoding a germline-specific Argonaute required for piRNA-mediated transposon repression (Williams and Rubin 2002), is among the top 200 genes with significantly higher expression in GSC-like cells (Fig. 2B, Supplemental Table 2).

**Fig. 2.**
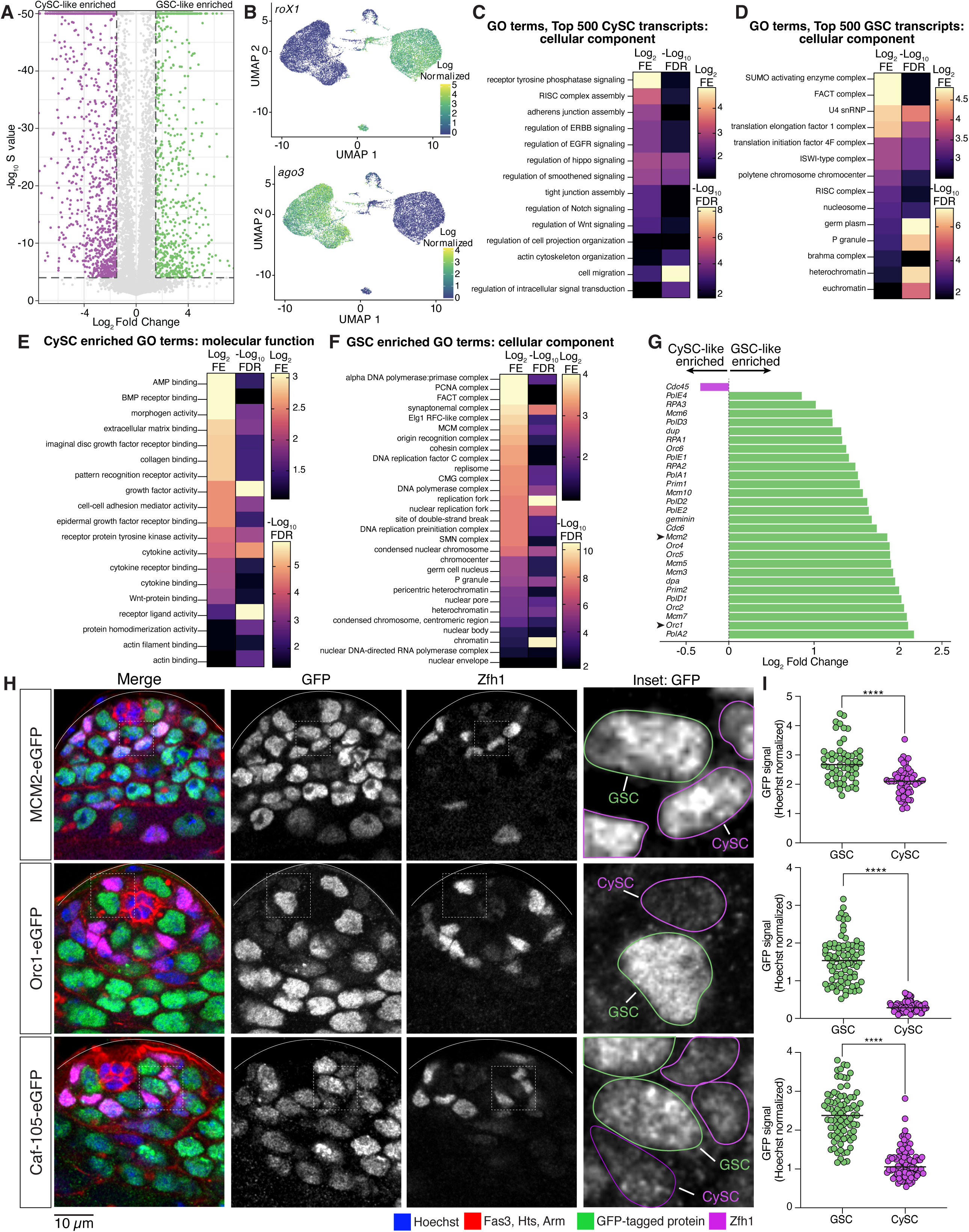

We performed Gene Ontology (GO) analysis to compare gene functions of expressed genes in GSC-like and CySC-like cells. We first analyzed the top 500 expressed transcripts based on their CPM (counts-per-million) values (top ∼2.8%). We chose this approach to complement our differential gene expression analysis to address the limitation that genes expressed at similar levels in both cell types would otherwise be undetected. In this way, we found that CySCs are enriched with transcripts encoding cell-cell adhesion molecules and signaling pathway components at high levels (Fig. 2C, Supplemental Table 3). For example, Cadherin-N (*cadN*) transcript level in the CySC-like cells (256.04 CPM) was higher than in GSC-like cells (0.17 CPM) (Supplemental Table 1). Components from several signaling pathways, including the ErbB, Epidermal Growth Factor (EGF), Hippo, Wnt, and Notch pathways, were identified as expressed highly in CySC-like cells (Fig. 2C, Supplemental Table 3), consistent with their roles in regulating CySC fate specification, division, and germline cyst enclosure (Schulz et al. 2002; Parrott et al. 2012; Kiger et al. 2000; Hudson et al. 2013; Ng et al. 2019). We also performed GO analysis on the 1,794 significantly higher differentially expressed transcripts in CySCs. These transcripts are also enriched for genes involved in intercellular communication, including paracrine signaling pathway components and adhesion proteins (Fig. 2E, Supplemental Fig. S3A, Supplemental Table 3). For example, non-cell-autonomous Bone Morphogenic Protein (BMP) signaling in somatic gonadal cells is essential for GSC maintenance and differentiation (Shivdasani and Ingham 2003; Leatherman and Dinardo 2010; Kawase et al. 2004). Components of this pathway, including the receptor Thickveins (Tkv) and the ligands Decapentaplegic (Dpp) and Glass bottom boat (Gbb), are significantly higher expressed in CySC-like cells in our analysis (Supplemental Table 2). Together, these profiles align with the known supportive roles of somatic gonadal cells in spermatogenesis.

In contrast to CySC-like cells, the GO analysis of transcripts in GSC-like cells indicates the top 500 genes expressed in GSC-like cells are primarily chromatin regulators, such as the Facilitates Chromatin Transcription (FACT), Imitation Switch (ISWI), and Brahma complexes (Fig. 2D, Supplemental Table 3) (Formosa and Winston 2020; Tyagi et al. 2016; Hu et al. 2021). For example, FACT complex subunits were expressed approximately five times higher in GSCs than in CySCs (Supplemental Table 1). We next examined the 827 significantly higher differentially expressed genes in the GSC-like cell cluster. The GSC-like transcriptome is differentially enriched for protein complexes involved in chromatin regulation, such as DNA replication, heterochromatin maintenance, and chromatin accessibility (Figs. 2F-G, Supplemental Figs. S3B-D, Table 1, Supplemental Table 3). Further, several chromatin-associated complexes contain multiple components, all significantly differentially expressed in GSC-like cells (Table 1). For example, genes encoding both FACT complex subunits, five Minichromosome Maintenance 2-7 (MCM2-7) complex components, and four Origin Recognition Complex (ORC) components, are all enriched in GSC-like cells compared to CySC-like cells (Table 1).

To control for potential biases in cell cycles, we normalized transcript representation across the G_1_, S, and G_2_/M cell cycle phases such that both cell types have equivalent percentages of cells in each stage (33% G_1_, 33% S, 33% G_2_/M, Supplemental Fig. S3F). This adjustment did not alter the enrichment pattern of genes, confirming that elevated replication gene expression in GSC-like cells reflects a true biological difference rather than differences in cell cycle phase distribution.

We next tested whether the enrichment of DNA replication factors detected in GSC-like cells from the *upd-*tumor also occurs in GSCs from non-tumor wild-type testes. Using genome editing, we generated knock-in fly lines with *gfp*-tagged endogenous *mcm2*, *orc1*, and *caf-p105*, genes considered differentially enriched in GSC-like cells (Fig. 2G, Supplemental Fig. S3D, Table 1). We then performed immunostaining on non-tumor testes to examine the expression patterns of these GFP-fusion proteins and quantified GFP signals in wild-type GSCs and CySCs (Figs. 2H-I, Supplemental Fig. S3G). In all instances, normalized GFP signal was significantly higher in GSCs compared to CySCs (Fig. 2I), demonstrating that the protein levels in wild-type testes recapitulate the transcript enrichment identified in *upd-*tumor samples. Together, these scRNA-seq data support the usage of *upd-*tumor testes in uncovering key characteristics of wild-type stem cells through genomic strategies.

### Cell type-specific chromatin landscapes profiled by H3K4me3, H3K27me3, and H3K9me3

The high expression of chromatin regulators in GSC-like cells, revealed by our scRNA-seq analysis, suggests their potential roles in shaping distinct chromatin landscapes to regulate germline functions. This prompted us to investigate cell-type-specific chromatin patterns in GSC-like and CySC-like cells, respectively. To achieve this, we optimized a low-input cell-type-specific chromatin profiling method using *upd-*tumor testes, which employs micrococcal nuclease (MNase) digestion of targeted chromatin (Fig. 3A). In this method, we co-expressed the Upd ligand and histone H3-GFP fusion protein in either GSC-like cells (*nanos*>*upd*, *H3-GFP*) or CySC-like cells (*tj*>*upd*, *H3-GFP*), as described earlier. MNase digestion specifically targeted chromatin containing H3-GFP using an anti-GFP antibody (Fig. 3A). After separating the soluble (GFP-containing) from insoluble (GFP-negative) chromatin, 5% of the soluble chromatin was sequenced as ‘input’, while the remaining chromatin was prepared for low input ChIP-sequencing (ChIP-seq) using antibodies against H3K4me3, H3K27me3, or H3K9me3 (Fig. 3A). We chose these histone modifications because they represent distinct chromatin domains: H3K4me3 (euchromatin), H3K27me3 (facultative heterochromatin), and H3K9me3 (constitutive heterochromatin) (Kouzarides 2007).

**Fig. 3.**
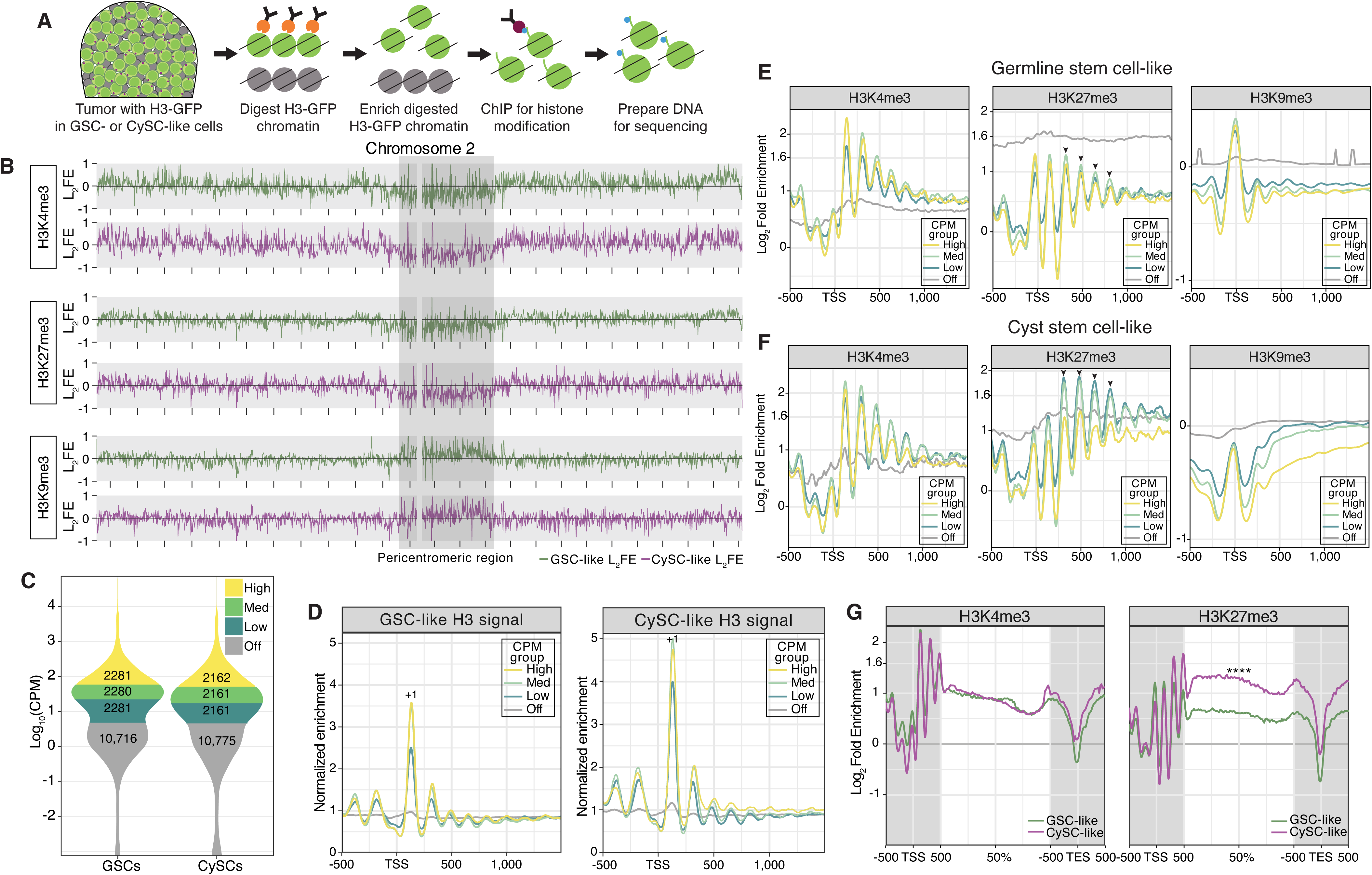

In agreement with previous findings (Kharchenko et al. 2011), H3K4me3 and H3K27me3 are enriched along the euchromatic chromosome arms but depleted in the pericentromeric regions of Chromosomes 2, 3, and X (Fig. 3B, Supplemental Fig. S4A). In contrast, H3K9me3, which associates with constitutive heterochromatin, shows enrichment at pericentromeric regions (Fig. 3B, Supplemental Fig. S4A). Together, these results demonstrate that our cell-type-specific chromatin profiling assay accurately recapitulates chromatin signatures consistent with the known distributions of H3K4me3, H3K27me3, and H3K9me3 (Kharchenko et al. 2011).

To investigate relationships between chromatin patterns and cell-type-specific gene expression, we integrated our cell-type-specific chromatin profiles with the scRNA-seq results. Focusing on the GSC-like and CySC-like clusters, we defined a transcript as expressed (‘on’) if its CPM was above 5, a threshold that reliably separates known cell-type-specifically expressed genes from non-expressed genes (CPM< 5, ‘off’). For example, using this cutoff, *ago3* is classified as ‘off’ in the CySC-like cluster (CPM = 2.11), but is robustly expressed in the GSC-like cluster (CPM = 697.21) (Fig. 2B, Supplemental Table 1), indicating strong cell type-specific expression (Williams and Rubin 2002). We further classified ‘on’ genes into three categories based on their transcript levels: low, medium, and high (Fig. 3C). The GSC-like and CySC-like clusters contained similar numbers of ‘on’ and ‘off’ genes (GSCs: 6,842 ‘on’ and 10,716 ‘off’; CySCs: 6,484 ‘on’ and 10,775 ‘off’), while the remaining three smaller clusters displayed comparable numbers of ‘on’ and ‘off’ genes (Supplemental Fig. S4B).

Using these categories, we plotted mononucleosomal enrichment to understand how nucleosomal position and density influence gene expression. The phased distribution of the H3 signal, indicative of nucleosome positioning, displays the characteristic mononucleosomal pattern generated by the MNase (Fig. 3D). This approach reveals the strongly positioned +1 nucleosome near the transcription start sites (TSSs) of actively expressed genes in both cell types, with positioning strength diminishing as gene expression levels decrease (Fig. 3D, Supplemental Table 4). This observation aligns with the known association between strongly positioned nucleosomes and chromatin organization at active genes, underscoring the role of nucleosome positioning in regulating gene expression and further validating our experimental approach (Chereji et al. 2018).

### H3K27me3 plays a more prominent role in CySC-like cells compared to GSC-like cells

To investigate the relationship between histone modifications and gene expression, we analyzed enrichment of H3K4me3, H3K27me3, and H3K9me3 at TSS relative to transcript levels using input-normalized Log_2_ Fold Enrichment (L_2_FE). In both GSC-like and CySC-like cells, H3K4me3 enrichment positively correlates with gene expression (Supplemental Figs. S4C, S4E). Consistently, average enrichment profiles show a pronounced H3K4me3 peak at the +1 nucleosome of expressed genes (Figs. 3E-F, Supplemental Table 4). In contrast, silenced genes display higher H3K9me3 enrichment than expressed genes (Figs. 3E-F, Supplemental Figs. S4C, S4E), with transposable elements having particularly strong enrichment, consistent with the role of H3K9me3 in marking constitutively repressed regions (Supplemental Fig. S4D, Supplemental Table 4). Because gene expression categories were defined independently for each cell type, we also compared average gene profiles of genes expressed in both cell types and uniquely in one *versus* the other (Supplemental Fig. S4F, S4G). The overall patterns of individual histone modifications were similar across gene sets, regardless of cell type, indicated that no single histone mark alone can predict cell-specific gene expression (Supplemental Fig. S4G).

Among the three profiled histone modifications, H3K27me3 displays distinguishable cell-type-specific differences between GSC-like and CySC-like cells, indicating potentially distinct roles for this modification in these two cell types (Supplemental Figs. S4C, S4E). In GSC-like cells, H3K27me3 shows moderate enrichment across all expressed gene categories (Fig. 3E, Supplemental Figs. S4C, S4E, S4G), whereas silenced genes display stronger enrichment despite lacking well-positioned nucleosomes (Fig. 3E). This non-canonical H3K27me3 enrichment resembles the pattern observed in female *Drosophila* GSCs, where H3K27me3 is detectable at both inactive and active gene loci (Deluca et al. 2020).

In contrast, in CySC-like cells, H3K27me3 is most enriched at the lowest-expressed genes and gradually decreases with increasing gene expression levels (Fig. 3F, Supplemental Figs. S4C, S4E). This H3K27me3 pattern is consistent with its well-established role in gene repression. Further, in CySC-like cells, H3K27me3 enrichment increases at the +3 nucleosome position and extends into the gene body for medium- and low-expression genes, a distinct feature not observed in GSC-like cells (Figs. 3E-F).

Previous studies demonstrate that broad H3K4me3 domains mark genes critical for cell identity and function in stem and progenitor cells. As cells differentiate and lose potency, these broad H3K4me3 domains become refined (Zhang et al. 2021). Further, broad H3K27me3 enrichment across gene bodies is associated with gene repression (Young et al. 2011). To investigate whether broad histone modification patterns are present in the two populations of stem cells in testis, we analyzed histone modification enrichment across gene bodies in genes expressed in both GSC-like and CySC-like cells (Fig. 3G). Our analysis revealed similar H3K4me3 enrichment along gene bodies in both cell types (Fig.3G, Supplemental Table 4). However, H3K27me3 was significantly enriched along gene bodies in CySC-like cells (Fig. 3G, Supplemental Table 4). This differential enrichment indicates a cell-type-specific regulatory mechanism for which H3K27me3 plays a more prominent role in CySC-like cells than in GSC-like cells.

Previous functional studies in *Drosophila* testes demonstrate that the histone methyltransferase Enhancer of Zeste [E(z)], which generates H3K27me3, plays a critical non-cell autonomous role in the CySC lineage by preventing ectopic expression of a key somatic transcription factor in germ cells (Eun et al. 2014). Conversely, inactivation of E(z) in the germline results in only mild defects, which arise in later-stage germ cells rather than GSCs (Eun et al. 2017). This indicates H3K27me3 may not play a large role in the early germline. Consistent with these findings, our scRNA-seq data show moderate *E(z)* expression in both GSC-like (CPM = 74.33) and CySC-like (CPM = 20.89) cells (Supplemental Fig. S4H, Supplemental Table 1). However, transcript levels for *Utx*, the demethylase responsible for erasing the H3K27me3 modification, are more highly expressed in the GSC-like cluster (GSC-like CPM = 64.17, CySC-like CPM = 31.06) (Supplemental Fig. S4I, Supplemental Table 1). Together, these findings highlight a role for E(z) and H3K27me3 in the CySC lineage, where they regulate the identity and activity of the GSC lineage in a non-cell-autonomous manner. Moreover, these data suggest a non-canonical H3K27me3 chromatin pattern in male GSC-like cells, resembling the pattern detected in female GSCs (Deluca et al. 2020).

### Distinct replication timing profiles in GSC-like and CySC-like cells

Chromatin organization plays an integral role in regulating transcription and DNA replication. Given the differences in chromatin landscapes between GSC-like and CySC-like cells, we aimed to understand how replication timing differs between these cell types using Replication sequencing (Repli-seq) (Fig. 4A, Supplemental Figs. S5A-C) (Marchal et al. 2018). First, we dissected *upd*-tumors expressing H3-GFP in either GSC-like or CySC-like cells and incubated them with the thymidine analog Bromodeoxyuridine (BrdU) to label replicating DNA. Next, GSC-like or CySC-like nuclei were extracted and sorted *via* flow cytometry into four S-phase fractions corresponding to early, early-mid, late-mid, and late S-phase, based on their DNA content (Fig. 4A, Supplemental Figs. S5A-C). BrdU-labeled DNA was then immunoprecipitated, followed by library preparation and sequencing (Fig. 4A).

**Fig. 4.**
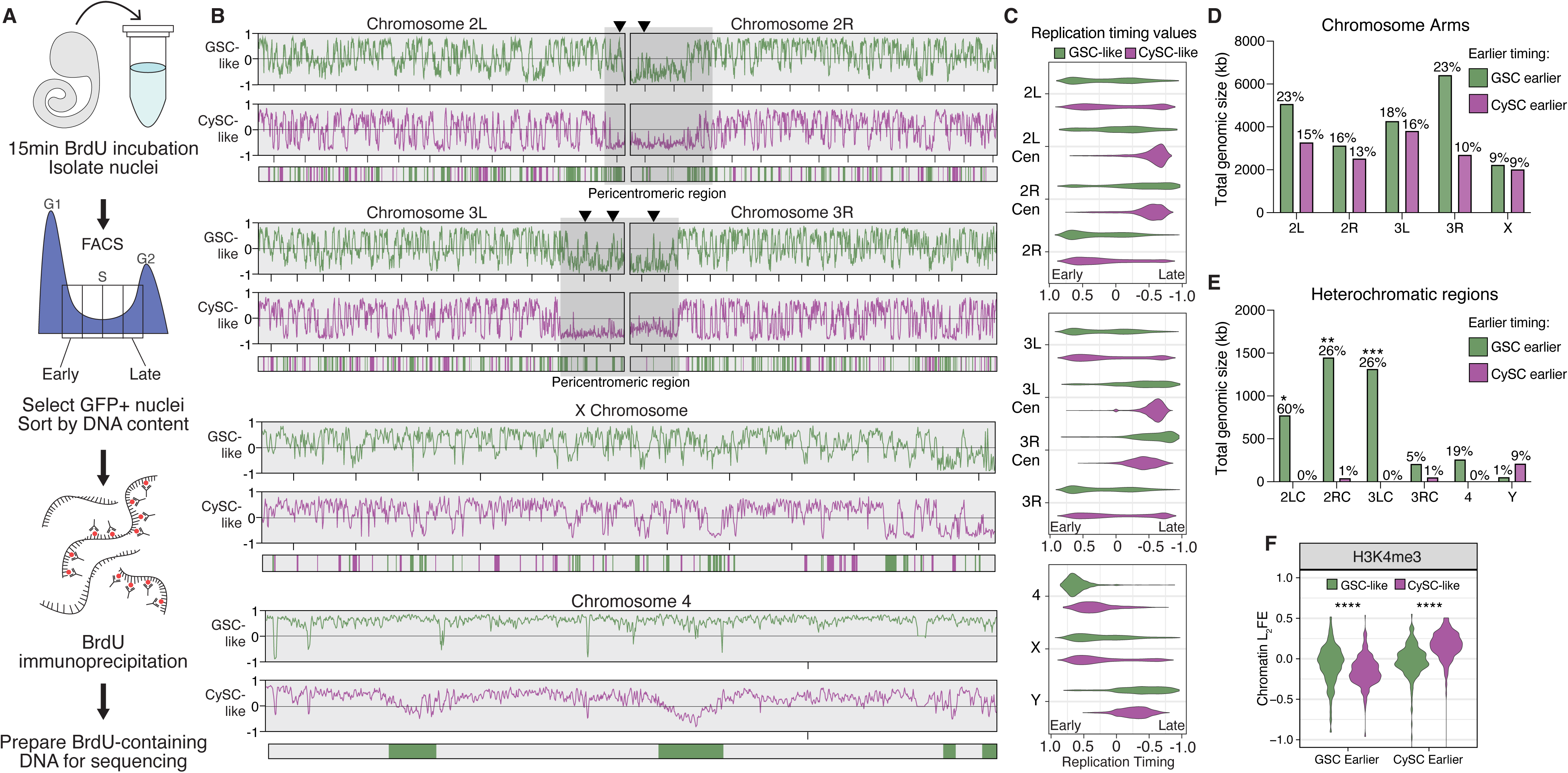

We generated replication timing profiles to identify early and late replicating genomic regions on each chromosome. In these profiles, enrichment of Repli-seq reads in the early fraction yields a positive score, just as late fraction enrichment yields a negative score. Consistent with known replication timing patterns, pericentromeric heterochromatin is largely late-replicating in both GSC-like and CySC-like cells (Figs. 4B-C). On the other hand, in both cell types, the chromosomal arms of Chromosomes 2, 3, and the X replicate throughout S-phase, with some regions replicating early and others late (Figs. 4B-C). The 2^nd^ and 3^rd^ Chromosomes comprise 70-80% of the *Drosophila* genome and display distinct chromatin landscapes between the chromosome arms and pericentromeric regions (Fig. 3B, Supplemental Fig. S4A). To examine replication progression relative to these chromatin domains, we plotted replication timing values for these two major autosomes, separating the arms from the pericentromeric regions. Minimal replication timing differences were detected between GSC-like and CySC-like cells along the major autosomal arms (Fig. 4C). Notably, while pericentromeric regions for both major autosomes were consistently late-replicating in CySC-like cells, they displayed early replication features in GSC-like cells (Figs. 4B-C, Supplemental Table 5). The 4^th^ Chromosome replicates early in GSC-like cells, despite its known heterochromatic nature (Figs. 4B-C) (Haynes et al. 2007). Overall, GSC-like cells exhibited a trend where traditionally heterochromatic regions replicate earlier than expected, suggesting previously unrecognized roles for these regions in regulating GSC functions.

We next compared replication timing values between GSC-like and CySC-like cells to identify genomic regions with significantly different timing regimes. Using 3kb sliding windows, we tested the null hypothesis that replication timing is equal between the two stem cell types. Nested peak calling was then performed to identify regions at least 20kb in size with significant timing differences between the two cell types. Regions were then categorized as ‘GSC earlier’ or ‘CySC earlier’ depending on whether they replicate earlier in GSC-like or CySC-like cells (Fig. 4B, Supplemental Fig. 5D). Overall, 25.12 Mb of the genome replicates earlier in GSC-like cells compared to CySC-like cells, while 14.588 Mb replicates earlier in CySCs-like cells (Fig. 4D-E, Supplemental Table 6). In addition to total genomic size, we calculated the fraction of each chromosome that replicates earlier to gain perspective on the scale of these differences (Supplemental Table 6). The total genomic size of earlier replicating regions is similar between GSC-like and CySC-like cells across the major autosomal arms and the X Chromosome (Fig. 4D). A notable exception is the right arm of Chromosome 3, where the total amount of kb replicating earlier in GSC-like cells is much greater than CySC-like cells. Accordingly, 23% of Chromosome 3R replicates earlier in GSC-like cells, whereas just 10% of this chromosome replicates earlier in CySC-like cells (Fig. 4D, Supplemental Table 6).

In heterochromatic regions, GSC-like cells exhibited a greater total genomic size of earlier-replicating DNA compared with CySC-like cells (Fig. 4E). These differences span substantial portions of the genome. For example, earlier-replicating domains in GSC-like cells account for 60% of the pericentromeric heterochromatin on the 2^nd^ Chromosome. A notable exception is the Y Chromosome, where 9% of the Y Chromosome replicates earlier in CySC-like cells than in GSC-like cells (Fig. 4E, Supplemental Fig. 5D). To examine whether differences in replication timing align with differences in histone modification enrichment, we analyzed L_2_FE values from our cell type-specific histone modification data. As expected, regions that replicate earlier in GSC-like cells show significantly greater H3K4me3 enrichment in this cell type (Fig. 4F, Supplemental Table 5). Similarly, earlier replicating regions in CySC-like cells are enriched for H3K4me3 in these cells (Fig. 4F, Supplemental Table 5).

These results indicate that cell type-specific replication timing differences align with cell type-specific chromatin features. For the first time, our results enable precise identification of specific chromosomal regions that differ in replication timing and chromatin landscapes between germline and somatic stem cell populations in the *in vivo* context.

### Replication timing of GSC-like cells is distinct from that of CySC-like and cultured *Drosophila* cells

While most studies on replication timing have focused on cultured cell lines, studies using *in vivo* cells from a normal developmental context revealed differences in replication timing profiles during cell fate specification. This highlights how dynamic variations in replication programs associated with changes in cellular potency cannot be captured by studying cultured cells alone (Armstrong et al. 2018; Pratto et al. 2021; Takahashi et al. 2024; Das et al. 2021; Siefert et al. 2017; Nakatani et al. 2023). To gain cell-type specific insights and compare replication timing between *in vivo* and *in vitro* cells, we compared our datasets with published replication timing data from two well-studied *Drosophila* cell lines, Kc167 and S2 (Lubelsky et al. 2014). Our analysis suggests that the replication timing profile of GSC-like cells is distinct from the other three cell types, while the two cell lines cluster together (Fig. 5A).

**Fig. 5.**
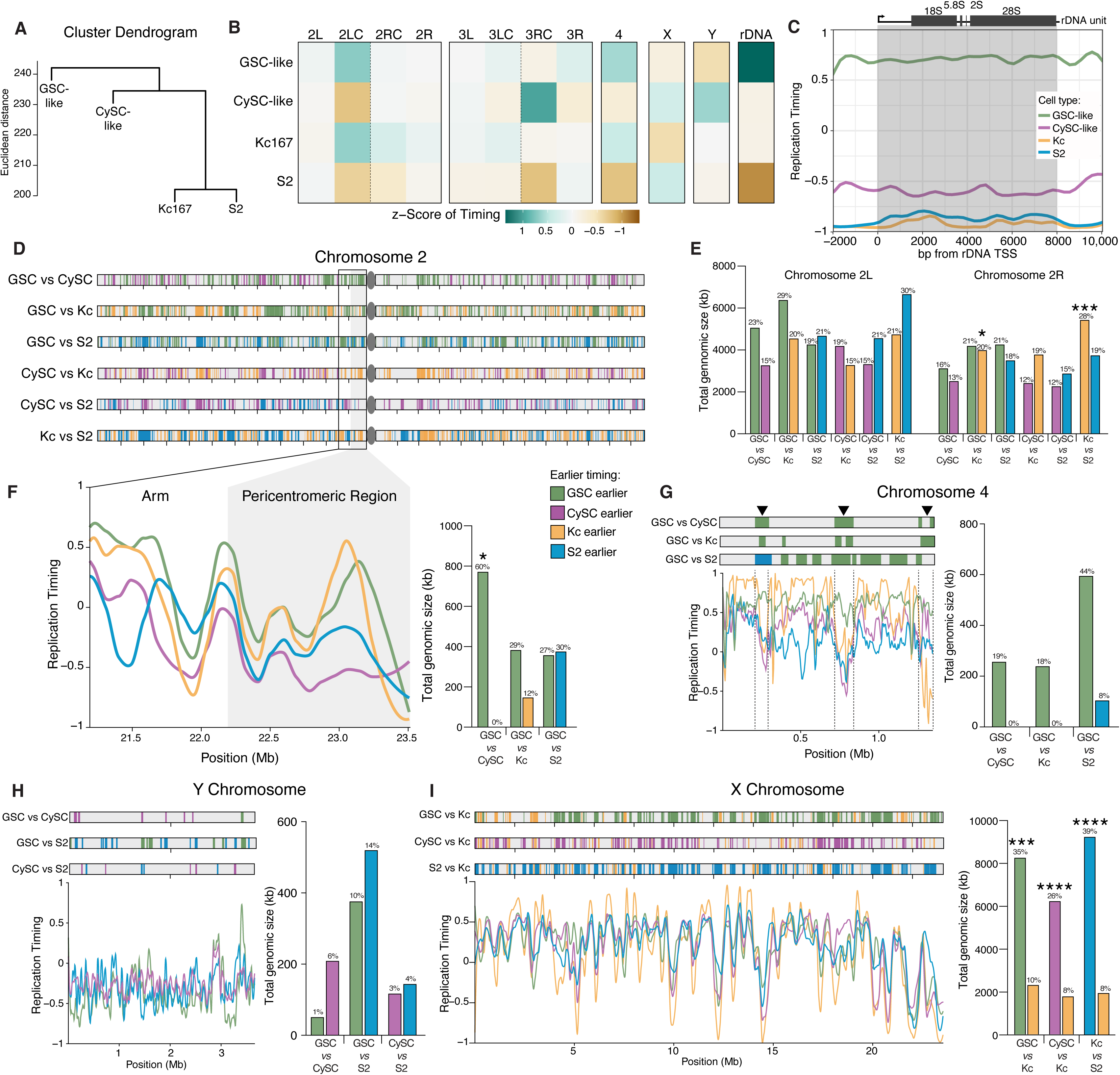

To facilitate cross-cell-type comparisons, we standardized replication timing along the reference genome to have unit variance, yielding RT *Z*-scores. We then categorized the genome into 12 chromosomal domains with their mean RT *Z*-scores: two major autosomal arms and their pericentromeric regions, the 4th, X, and Y Chromosomes, and a gene consensus for the ribosomal DNA (rDNA) locus (Fig. 5B). This allowed us to characterize negatively and positively correlated cell type profiles (Supplemental Fig. S6A). Among the four cell types, GSC-like and Kc167 cells show the strongest correlation in their chromosomal region RT *Z*-score profiles (Supplemental Fig. S6A). This is intriguing because Kc167 cells are female (XX karyotype) (Cherbas and Gong 2014). Similarly, CySC-like and S2 cells display positively correlated RT *Z*-scores, likely due to their shared somatic identity and male (XY) karyotype (Supplemental Fig. S6A).

We additionally summarized the chromosomal region RT *Z*-scores by their arithmetic mean (Fig. 5B). These summarized *Z*-scores, relative to the combined dataset mean, indicate how each genomic region in each cell type deviates from the dataset’s average (Fig. 5B). While chromosomal arms display no overall trends, this method identified earlier replication timing in GSC-like cells for the pericentromeric region of Chromosome 2L (Early-Mid), the 4th Chromosome (Early), and the rDNA locus (Early) (Fig. 5B). Although replication timing for the X Chromosome was found to be Early-Mid and the Y Chromosome to be Late-Mid in both GSC-like and CySC-like cells (Supplemental Table 5), the *Z*-scores revealed more genomic regions on both sex chromosomes that replicate earlier in CySC-like cells (Fig. 5B). Intrigued by the early replicating result at the rDNA locus, we plotted replication timing scores for each cell type across the 8kb transcriptional unit within the rDNA locus in the dm6 genome. Indeed, while the negative replication timing values for CySC-like, Kc167, and S2 cells indicate late replication, positive values in GSC-like cells demonstrate their cell-type-specific early replication timing (Fig. 5C).

Pairwise comparisons of replication timing profiles across the four cell types identified genomic windows with distinct replication timing. As shown above, we quantified the total size (kb) of earlier-replicating regions and mapped their genomic locations by cell type. Additionally, we calculated the corresponding proportion of each chromosomal domain that replicates earlier. The arms of the 2nd and 3rd Chromosomes show minimal differences in the amount, proportion, or location of cell-type-specific earlier-replicating regions (Fig. 5D-E, Supplemental Figs. S6B-C). Thus, differences in replication timing are present across the chromosome arms and collectively encompass 12-30% of these chromosome domains.

We next focused on regions where the mean RT *Z*-score suggests earlier replication in GSC-like cells, such as the pericentromeric region of Chromosome 2L and the 4^th^ Chromosome. In both cases, the total genomic size of earlier replicating regions is greater in GSC-like cells when compared with other cell types (Fig. 5F-G, Supplemental Figs. S6D-E, Supplemental Table 6). The pericentromeric region does not have specific locations with significantly different replication timing between the GSC-like cells and other cell types; rather, the entire region displays earlier replication timing in GSC-like cells than both CySC-like and S2 cells (Fig. 5F). Kc167 and GSC-like cells have comparable replication timing patterns within this region (Fig. 5F). In contrast to the pericentromeric region of Chromosome 2L, the 4th Chromosome contains three distinct regions, accounting for ∼19% of this chromosome, with comparatively earlier replication timing in GSC-like cells (Fig. 5G, Supplemental Fig. S6E).

The Y Chromosome generally replicates late in S-phase across all three male cell types (Fig. 5H). However, CySC-like cells show a relatively earlier replication timing profile, as reflected by their mean RT *Z*-score (Fig. 5B). Indeed, the total genomic size of earlier-replicating regions is larger in both CySC-like and S2 cells compared to GSC-like cells, while CySC-like and S2 cells show similar values (Fig. 5H, Supplemental Table 6). Unlike the 4th Chromosome, replication timing for the Y Chromosome does not reveal any consistent earlier-replicating regions and these domains compose small fractions of the Y Chromosome (Fig. 5H). However, a few specific loci show different replication timing when comparing GSC-like to CySC-like cells, which are subject for subsequent analyses (Fig. 5H).

Finally, when comparing male cells (GSC-like, CySC-like, or S2) to female Kc167 cells, the X Chromosome consistently replicates earlier in the male cell, whereas the average replication timing of the X Chromosome was similar among male cell types (all with a summary RT value of 0.48 ± 0.01) (Fig. 5I, Supplemental Fig. S6F, Supplemental Table 5). This predominance of early-replication of the X Chromosome in male cells likely reflects the accessible chromatin environment that facilitates male-specific X Chromosome dosage compensation (Urban et al. 2017). Early X Chromosome replication is preserved in GSC-like cells, despite the absence of dosage compensation in this cell type (Fig. 2D) (Meiklejohn et al. 2011). Our scRNA-seq further supports this note, showing no evidence of X-linked gene dosage compensation in GSC-like cells or primary spermatocytes, based on the X-to-autosome (X:A) transcript abundance ratios (Supplemental Fig. S6G). Overall, our analysis identifies key genomic regions with cell type-specific replication timing differences, including heterochromatic pericentromeric regions and the 4th, X, and Y Chromosomes, suggesting distinct functional requirements for these domains across different cell types.

### Cell-type-specific replication timing correlates with histone modification patterns

The temporal order of genome duplication in eukaryotic cells is intricately linked to chromatin structure and gene expression, both of which are cell-type-specific (Rhind and Gilbert 2013). To investigate this relationship, we integrated our replication timing data with the cell-type-specific histone modification profiles generated in this study. Additionally, we reanalyzed published Repli-seq and ChIP-chip data from Kc167 and S2 cells to compare those *in vitro* datasets with our *in vivo* results (Kharchenko et al. 2011; Lubelsky et al. 2014).

Consistent with previous findings in Kc167 and S2 cells (Eaton et al. 2011), H3K4me3 is enriched at early replicating regions, coinciding with genes that have the highest expression levels (Fig. 6A-B, Supplemental Figs. S7A-C). Accordingly, elevated H3K4me3 is present at the TSSs of genes located in the early-replicating fraction in both GSC-like and CySC-like cells (Supplemental Figs. S7E- F). In contrast, H3K9me3 is found in gene-poor and late-replicating regions across all cell types (Figs. 6A-B, Supplemental Figs. S7A-B, S7D-F).

**Fig. 6.**
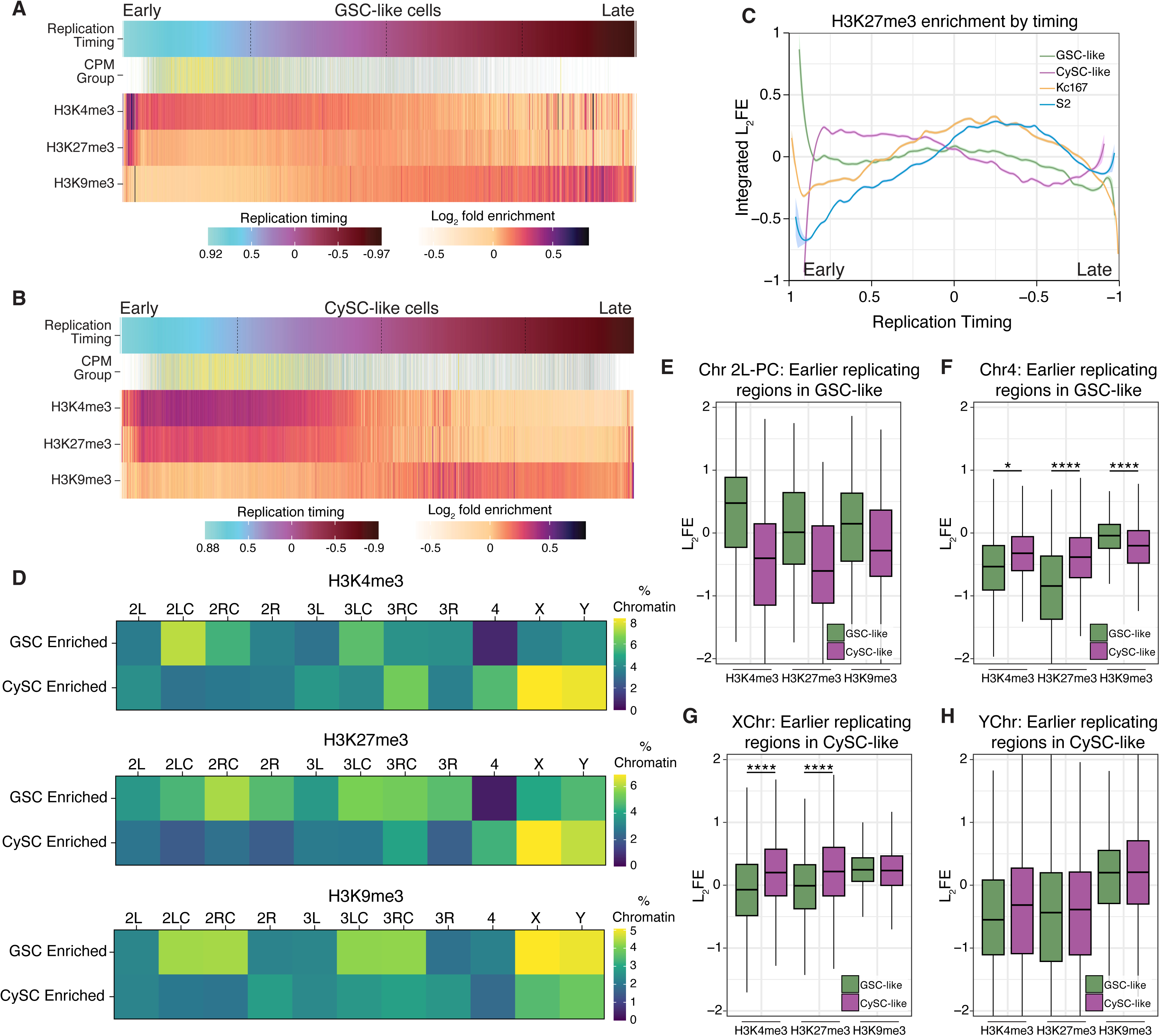

On the other hand, H3K27me3 displays distinct enrichment patterns at different S-phase stages across cell types. In CySC-like cells, H3K27me3 is present throughout early and early-mid S-phase (Figs. 6B-C), overlapping with H3K4me3, and is enriched at TSSs of genes replicating in these stages (Supplemental Fig. S7F). Notably, this H3K27me3 pattern is distinct from the other three cell types, further supporting the observation that there is a unique H3K27me3 chromatin signature in CySC-like cells. In contrast, H3K27me3 is predominantly enriched during mid S-phase in GSC-like, Kc167, and S2 cells (Figs. 6A, 6C, Supplemental Figs. S7A-B). Consistently, in GSC-like cells, H3K27me3 is most enriched at TSSs of genes replicated during early-mid S-phase (Supplemental Fig. S7E).

Our replication timing analysis revealed cell-type-specific timing regimes for pericentromeric heterochromatin on Chromosome 2L and globally for the 4th, X, and Y Chromosomes (Fig. 5). We next tested whether these regions have corresponding differences in chromatin landscapes. To compare histone modification differences across genomic regions between GSC-like and CySC-like cells, we defined a given locus as having cell-type-specific histone modification enrichment if its L_2_FE was >0.2 in one cell type and <-0.2 in the other. This analysis revealed strong overall correlations, suggesting broadly similar histone modification landscapes (Supplemental Fig. S7G). However, distinct GSC- or CySC-enriched histone modification regimes were also identified (Fig. 6D, Supplemental Fig. S7G, green and purple boxes). To quantify cell-type-specific chromatin features, we calculated the percentage of specific genomic regions enriched with each histone modification (Fig. 6D). For example, 7.4% of the pericentromeric region of Chromosome 2L in GSC-like cells is enriched for H3K4me3, compared to only 3.2% in CySC-like cells (Fig. 6D, Supplemental Table 7), aligning with its earlier replication timing in GSC-like cells (Fig. 5C). In contrast, the 4th Chromosome in GSC-like cells shows almost no specific enrichment of H3K4me3 or H3K27me3 (Fig. 6D, Supplemental Table 7), despite its markedly early replication in this cell type. In CySC-like cells, a greater percentage of chromatin on the X and Y Chromosomes is enriched with both H3K4me3 and H3K27me3 (Fig. 6D).

We next evaluated histone modification enrichment scores at regions with distinct replication timing to assess correlations between cell-type-specific replication timing and chromatin features. In GSC-like cells, earlier replicating regions in Chromosome 2L pericentromeric heterochromatin displayed higher H3K4me3 enrichment than in CySC-like cells (Fig. 6E, Supplemental Table 5). These earlier replicating regions on the 4th Chromosome in GSC-like cells do not display a similar increase in H3K4me3 L_2_FE values when compared with CySC-like cells (Fig. 6F, Supplemental Table 5), suggesting that the early replication of this chromosome may not be linked to a chromatin mark like H3K4me3.

In CySCs, the X and Y Chromosomes replicate earlier (Fig. 5C), and their earlier-replicating regions show higher levels of H3K4me3 and H3K27me3 compared to those in GSC-like cells (Figs. 6G- H, Supplemental Table 5). A similar observation was made for regions on the X and Y with earlier- replication in GSC-like cells (Supplemental Figs. S7H-I, Supplemental Table 5), indicating a chromosome- and cell-type-specific relationship between replication timing and histone modifications. In summary, this integrative analysis reveals a relationship between cell type- and chromosome-specific chromatin landscapes and DNA replication programs in *Drosophila* testis stem cells.

### Selected features within differentially replicating regions in germline and somatic lineages

To explore the relationship between cell-type-specific replication timing and gene expression, we assessed gene-specific replication timing and expression differences between GSC-like and CySC-like cells within regions identified as differentially replicating. In the pericentromeric region of Chromosome 2L, several differentially expressed genes, such as *Marf1*, *mir-2490*, and *CR46253*, replicate earlier in GSC-like cells than in CySC-like cells (Figs. 7A-B). While *Marf1* represses *nanos* expression and promotes meiotic progression during oogenesis in the female germline (Kawaguchi et al. 2020), the functions of *Marf1*, *mir-2490,* and *CR46253* in the male germline remain unknown. Notably, the genomic loci for these regulatory RNAs are associated with Orc2 binding sites, suggesting potential replication origins within this region (Fig. 7B). These observed differences in replication timing between GSC-like and CySC-like cells may reflect cell-type-specific origin usage that could arise from transcriptional differences within this region.

**Fig. 7.**
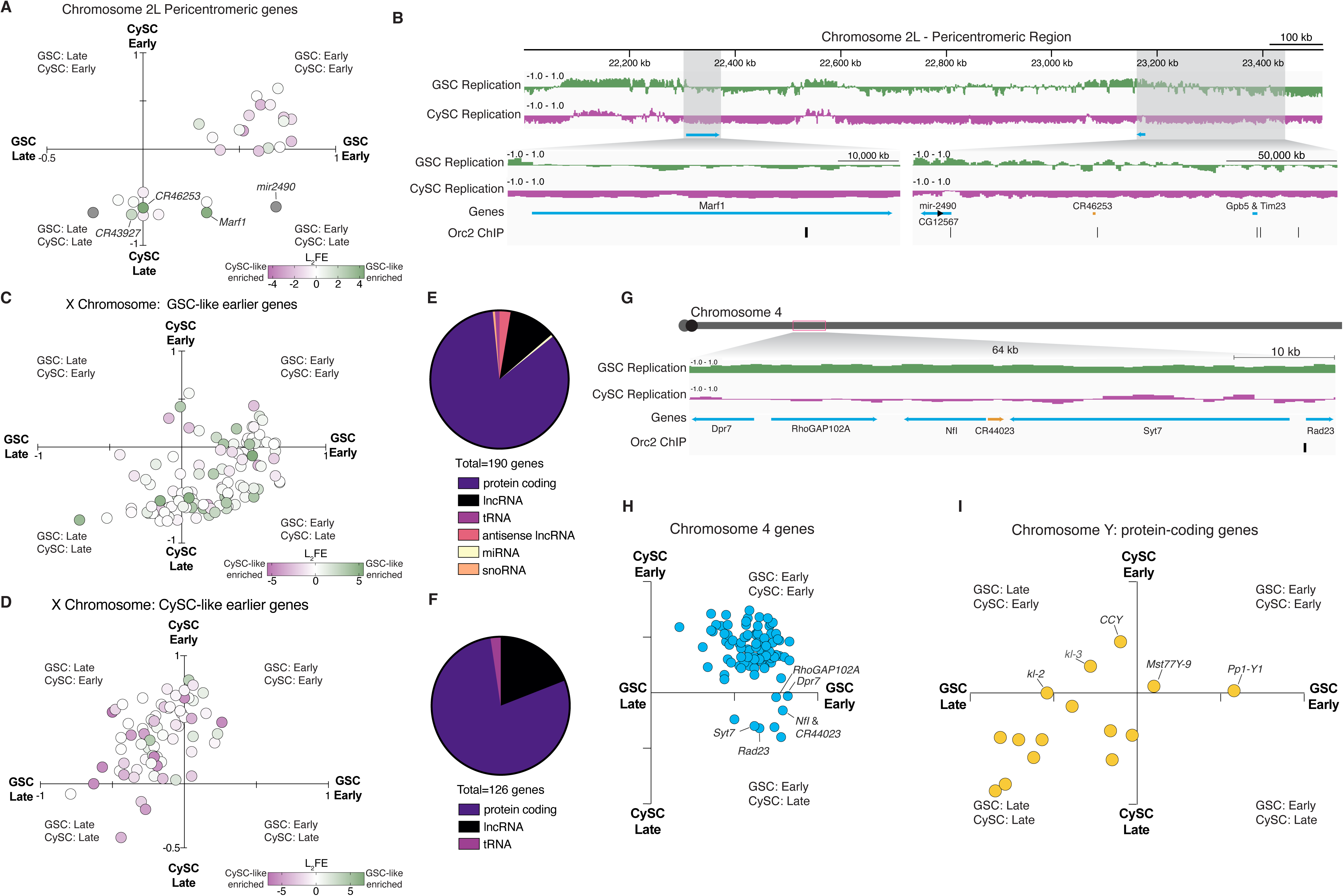

Extending beyond Chromosome 2L, the X Chromosome also exhibits distinct replication timing and histone modification enrichment in a cell-type-specific manner. In male cells, the X Chromosome generally replicates early, with CySC-like cells displaying higher H3K4me3 enrichment (Fig. 6D). The correlation between early replicating regions and active histone modification enrichment further supports the connection between chromatin state and replication timing. Categorizing X-linked genes by cell-type-specific replication timing revealed a strong alignment between earlier replication and higher expression in the corresponding cell type (Figs. 7C-D). Notably, GSC-like cells showed more non-coding genes in early-replicating regions, suggesting a potential regulatory role for non-coding RNAs in this cell type (Figs. 7E-F).

Similarly, distinct replication patterns are also observed on the 4th Chromosome. Several loci on the 4th Chromosome replicate earlier in GSC-like cells across all pair-wise comparisons (Fig. 5G). One 64kb domain contains six tandem genes that synchronously replicate early (Fig. 7G), clustering in the Early GSC, Late CySC quadrant (Fig. 7H). This region has five protein-coding genes and one uncharacterized long non-coding RNA. This region is also associated with an Orc2 binding site, suggesting cell-type-specific origin usage (Fig. 7G).

The Y Chromosome also follows distinct replication timing patterns, with certain genes exhibiting early replication in a cell-type-specific manner despite its largely late-replicating nature. Functional data on the 16 annotated protein-coding genes on the Y Chromosome remains limited. For example, the *CCY* gene replicates early in CySC-like cells (Fig. 7I) and is essential for sperm individualization during late spermiogenesis, though its role in somatic gonadal cells remains unknown (Hafezi et al. 2020; Zhang et al. 2020). The presence of these genes in cell-type-specific early-replicating regions suggests their unexplored roles within the somatic lineage.

Taken together, these findings reveal links between replication timing, chromatin state, and cell identity. In summary, our replication timing data aligns with the cell-type-specific chromatin landscape, highlighting distinct regulatory regimes for the X, Y, and 4th Chromosomes in the soma *versus* germline. A broader pattern emerges: regions that are typically heterochromatic in differentiated cells replicate earlier in GSC-like cells. This is reminiscent of findings in *Drosophila* female GSCs and cultured mouse embryonic stem cells, which also exhibit reduced heterochromatic marks compared to differentiated cells (Pang et al. 2023; Rausch et al. 2020). Thus, the male germline provides a valuable system to study gene expression, chromatin regulation, and DNA replication in endogenous adult stem cell systems of a multicellular organism.

## DISCUSSION

Understanding the mechanisms underlying cell fate specification and fate transitions is essential to developmental and stem cell biology. In this study, we integrated transcriptional profiling, chromatin landscape analysis, and one of the first *in vivo* replication timing studies in adult stem cells to identify both shared and distinct regulatory mechanisms of *Drosophila* male germline and somatic cyst stem cells. We further compared our *in vivo* datasets with *in vitro* cultured *Drosophila* cell lines. In this way, we uncovered stem cell-type-specific differences in genomic and epigenomic features. To enable these studies, we used a stem cell tumor model that provided sufficient material for multi-omic profiling. While our results were derived from this model, many of our findings align with known features of wild-type germline and cyst stem cells. This supports the model’s utility as a platform for generating hypothesis and for guiding future investigations in wild-type tissue.

Our transcriptional profiling highlights a fundamental regulatory dichotomy between GSC-like and CySC-like cells. First, CySC-like cells rely on intercellular communication, with many highly expressed genes associated with signaling pathways that coordinate germline-soma interactions (Vidaurre and Chen 2021; Papagiannouli 2014). Second, GSC-like cells likely have a greater focus on regulating the chromatin landscape, with an overrepresentation of DNA replication and chromatin-modifying factors.

This difference aligns with the well-established roles of intercellular signaling in coordinating GSC and CySC identity and function. While intercellular communication appears an essential function for the somatic lineage, the strong enrichment of chromatin factors in GSCs suggests that chromatin regulation plays a more dominant role in regulating germline stem cell identity and activity.

### Cell-type-specific heterochromatin replication timing

Our study reinforces the well-established relationship between chromatin features and replication timing, where transcriptionally active euchromatic regions replicate early and transcriptionally inactive heterochromatic regions replicate late. However, by comparing these dynamics between two distinct adult stem cell populations *in vivo*, we find stem cell-type-specific differences. Notably, we found that heterochromatic domains have locations that replicate earlier in GSC-like cells compared to CySC-like and cultured cells, suggesting fundamental differences in heterochromatin regulation between germline and somatic lineages.

A distinction between GSC-like and CySC-like cells is the abundance and broad distribution of H3K27me3, a facultative heterochromatin mark established by the Polycomb-complex that is typically associated with gene silencing and cell identity maintenance (Wiles and Selker 2016). In CySC-like cells, H3K27me3 enrichment displays a canonical pattern, where lower expressed genes exhibit higher levels of this mark. It is possible that the extensive presence of H3K27me3 in CySCs serves to restrict gene expression and reinforce the cyst cell fate. This may reflect their unique division pattern - a single asymmetric cell division produces a self-renewed CySC daughter and a differentiating cyst cell (CC) daughter. Two post-mitotic CCs form a unit to support the differentiating germline for the duration of spermatogenesis, which takes approximately 10 days at 25°C (Fabian and Brill 2012; Zoller and Schulz 2012). Therefore, the restriction of H3K27me3 could reinforce somatic cyst cell identity, reflecting an early commitment to a terminal fate in this lineage.

In the female *Drosophila* germline lineage, constitutive pericentromeric heterochromatin formation begins in the germline cyst progeny of GSCs, while Polycomb-mediated gene repression suppresses male germline and somatic lineage genes (Pang et al. 2023). Further genomic and genetic studies in the female germline revealed an open chromatin environment in GSCs (Deluca et al. 2020). In female GSCs, there is a non-canonical H3K27me3 distribution, whereby this modification is present in low amounts on active chromatin as well as on transcriptionally inactive loci (Deluca et al. 2020). Here, we observe a similar trend in male GSCs, where there is a roughly equivalent amount of H3K27me3 across all gene categories regardless of their expression levels, suggesting that low, non-gene-specific H3K27me3 may be an epigenetic feature of undifferentiated cells (Deluca et al. 2020). However, in testes enriched with mitotic differentiating spermatogonial cells, the H3K27me3 pattern displays the canonical inverse correlation between enrichment and gene expression levels (Gan et al. 2010). As germ cells progress to meiotic spermatocytes, Polycomb function is counteracted, likely to allow de-repression of sperm differentiation genes (Chen et al. 2005, 2011). Taken together, these findings indicate developmentally regulated Polycomb activity across distinct germ cell stages in the *Drosophila* male germline.

Our finding that pericentromeric heterochromatin replicates early in GSCs aligns with published studies in mouse embryonic stem cells, which demonstrate that pericentromeric heterochromatin shifts from early to late replicating upon differentiation (Rausch et al. 2020). An additional key insight from that work was the significantly higher proportion of unidirectional replication forks in mouse embryonic stem cells (around 13%) compared to differentiated human cell lines (around 7%) (Rausch et al. 2020).

Similarly, a notable 35% of replication forks are unidirectional in the *Drosophila* germline compared to ∼17% in somatic cells (Wooten et al. 2019; Davis et al. 2025). This trend of higher unidirectional fork usage in less differentiated states appears to be consistent across species.

In addition, activated origins in mouse embryonic stem cells are spaced at half the distance of that in somatic cells, indicating that stem cells activate twice as many origins of replication (Rausch et al. 2020). Although future studies in *Drosophila* germline stem cells are needed to determine whether the higher incidence of unidirectional replication forks corresponds to reduced inter-origin distances, the elevated expression of DNA replication-related genes supports a model of increased replication demand in these cells. Furthermore, globally reduced H3K27me3 may create a permissive chromatin environment for increased unidirectional replication forks and origin activation. This supports the idea that unique replication dynamics may underlie specialized genome regulation in stem cells. These features may reflect developmental constraints on cell fate transitions or represent lineage-specific adaptations. Notably, our observation of early replication at the ribosomal DNA (rDNA) locus, which is well-established to replicate unidirectionally (Gerber et al. 1997), supports this model.

In parallel with these replication features, the *Drosophila* germline exhibits a striking pattern of histone inheritance: old histones, synthesized in the previous cell cycle, are preferentially incorporated onto the leading strand during DNA synthesis, while newly synthesized histones are biased toward the lagging the strand (Wooten et al. 2019; Davis et al. 2025). This produces sister chromatids asymmetrically enriched with different histone populations, which are then differentially segregated such that old-histone-enriched sisters are retained in the self-renewing germline stem cell while the gonialblast inherits new-histone-enriched sister chromatids (Ranjan et al. 2019; Tran et al. 2012). The elevated incidence of unidirectional forks may facilitate this global histone asymmetry, enabling extended tracts of old or new histone deposition. Early unidirectional replication at the rDNA locus may serve as a key organizing feature in establishing such genome-wide histone asymmetry in *Drosophila* GSCs. Together, these mechanisms may ensure the faithful inheritance of epigenetic information during stem cell asymmetric cell division, by preserving stem cell identity while allowing for flexible gene regulation in the differentiating progeny.

Recent studies in budding yeast suggest rDNA replication can influence anaphase onset, with extended rDNA repeats potentially delaying replication to prevent premature chromosome segregation (Kwan et al. 2023). In *Drosophila* germ cells, rDNA copy number decreases during aging, yet can be restored in GSCs of young males in the following generation (Lu et al. 2018; Nelson et al. 2023; Watase et al. 2022). The mechanism underlying this generational recovery remains unknown. Early replication of the rDNA locus in GSC-like cells may present a strategy for maintaining or restoring rDNA copy number for subsequent generations, which might be less efficient if replication occurred later in S-phase. This preferential timing of rDNA replication in stem cells may therefore serve a critical function in long-term germline and organismal maintenance. Future studies can further clarify how this unique replication feature contributes to stem cell genome integrity and lineage continuity.

### Chromosome-specific replication timing profiles and regulatory roles for non-coding RNAs

Our work also uncovered chromosome-specific differences in histone modification enrichment and replication timing, particularly for the X and 4^th^ Chromosomes. Historically, the 4^th^ Chromosome is considered as entirely heterochromatic, based on its high percentage of repetitive elements, lack of recombination, and high levels of HP1 and H3K9me2 (Haynes et al. 2007; Riddle et al. 2008). Previous studies in *Drosophila* suggest that the 4^th^ Chromosome largely shares chromatin features with pericentric heterochromatin, although it also contains some euchromatic regions (Riddle et al. 2008; Haynes et al. 2007). Unexpectedly, we found this chromosome replicates earlier in GSC-like cells than in CySC-like and other somatic cells. The biological significance of its early replication remains unclear. One possibility is that early replication of the 4^th^ Chromosome in GSC-like cells reflects a regulatory role for this chromosome that is specific to GSCs. Alternatively, this timing pattern may result from its similar chromatin composition to pericentromeric heterochromatin, rather than a regulatory process. Further studies are required to determine whether the early replication timing of the 4^th^ Chromosome in the male GSC-like cells has biological relevance for genome stability or gene regulation.

Our work also identified regulatory non-coding RNAs enriched in genomic regions undergoing coordinated changes in replication timing, chromatin state, and transcriptional activity, including those within the 4^th^ Chromosome and the pericentromeric heterochromatin of Chromosome 2. Although regulatory RNAs have historically been challenging to study, comparative analyses like ours allow for identifying them for further functional characterization. Previous studies have shown that approximately 30% of *Drosophila* testis-specific long non-coding RNAs (lncRNAs) function during late spermiogenesis, regulating nuclear condensation, sperm individualization, and other terminal differentiation processes (Wen et al. 2016). However, their roles in early spermatogenesis, particularly in stem cells, remains poorly understood. Notably, several non-coding RNAs located in differentially replicating regions were also differentially expressed in GSC-like cells. It is possible that their transcription promotes an open chromatin environment, facilitating origin recognition and early replication. Rather than acting through their RNA products, these transcripts may instead provide a mechanism to achieve the elevated origin activation observed in stem cells. An intriguing possibility is that the inherent directionality of RNA transcription could influence coordinated replication origin activation and directionality of fork progression. By identifying candidate non-coding RNAs situated in regions with stem cell-specific replication timing, our findings provide a foundation for investigating their functions in stem cell maintenance, differentiation, and genome regulation.

Our findings reinforce our knowledge that the male X Chromosome maintains a unique chromatin environment that supports its well-established role in dosage compensation in somatic cells (Samata and Akhtar 2018). In CySC-like cells, the X Chromosome displays signatures of open chromatin consistent with this function, including H3K4me3 (this study), H4K16ac, and early replication (Rastelli and Kuroda 1998; Anderson et al. 2023). In GSC-like cells where dosage compensation is inactive, the X Chromosome still replicates early. Previous studies in male and female *Drosophila* cell lines (Clone8 and Kc cells, respectively) similarly demonstrated that the X Chromosome replicates entirely during early S-phase, even at genes enriched with H4K16ac that did not undergo gene dosage compensation (Schwaiger et al. 2009). Based on these findings, H4K16ac was proposed to associate more with early replication than with transcriptional activity. However, our data show that early replication persists in male GSCs even in the absence of this mark. Together, these results possibly suggest a connection between replication timing of the X Chromosome and chromatin accessibility, as opposed to replication timing being solely linked to transcriptional output or H4K16ac enrichment (Schwaiger et al. 2009). Our findings support a model that the X Chromosome’s early replication timing in the male germline is maintained through dosage compensation-independent mechanisms, likely involving chromatin accessibility features.

In conclusion, by integrating transcriptional, chromatin, and replication timing analyses, our data provide a comprehensive framework for understanding the unique genomic regulation of male germline and somatic stem cells. Altogether, our findings reveal distinct fundamental divergence in regulatory mechanisms, including heterochromatin replication timing programs, chromosome-specific epigenetic differences, and potentially novel roles for lncRNAs in stem cell genome organization. These findings provide new insights into stem-cell specific genome regulation and establish a foundation for future studies on the epigenetic and replication dynamics of *in vivo* adult stem cells.

## METHODS

### Fly husbandry

Fly stocks were maintained on standard molasses food (Bloomington recipe) at 25°C. A complete list of fly stocks and sources is provided in the Supplemental Material. For genomic experiments, strains with switchable dual-tagged histone cassettes were introduced to a background containing a *UAS-upd* transgene (kindly provided by Dr. Stephen Dinardo, University of Pennsylvania, USA) using standard fly genetics. Male progeny of the correct genotype were aged for ∼2 days post-eclosion before dissection and preparation for scRNA-seq, ChIC-ChIP-seq, or Repli-seq. For immunostaining experiments, young males (1-5 days old) were collected from fresh flips.

### Immunostaining and image quantification

We used standard immunostaining approaches to perform both whole mount and squash immunostaining of wild-type and *upd*-tumor testes. A detailed description of both approaches is included in the Supplemental Materials along with a complete list of all antibodies used in this study. For whole-mount staining, dissected testes were fixed, incubated with primary and secondary antibodies, counterstained with Hoechst, and imaged by confocal microscopy. For the squash method, *upd*-tumor testes were ruptured, flash-frozen, ethanol-fixed, and stained with antibodies using similar incubation and wash steps prior to imaging. GFP-tagged Orc1, MCM2, and Caf-p105 fusion proteins were quantified from whole-mounted wild-type GSCs and CySCs using Imaris image analysis, with GFP intensity normalized to DNA content by Hoechst signal. Normalized GFP values were log_2_ transformed, plotted, and compared using a Mann–Whitney *U* test.

### 10x Genomics single cell RNA sequencing

For single-cell RNA sequencing, tumor testes from *nano*s>Upd, H3-GFP or *tj*>Upd, H3-GFP flies were dissociated into single cell suspensions using TrypLE Express and collagenase, followed by sequential filtration and washing to remove debris. Cell viability and counts were assessed with Trypan blue prior to library preparation. Single-cell RNA-seq libraries were generated using the 10x Genomics Chromium Next GEM Single Cell 3’ Kit v3.1 according to the manufacturer’s protocol, pooled to 4 nM, and sequenced using an Illumina NovaSeq S1 flow cell at the Johns Hopkins Genomics Core.

### Cell-specific sequential Chromatin Immunocleavage – Chromatin Immunoprecipitation (ChIC-ChIP) sequencing

Cell-specific sequential ChIC-ChIP was modified from an existing ChIC protocol optimized in the *upd-*tumor (Chandrasekhara et al. 2023). In this study, ChIC-ChIP was performed to profile the H3K4me3, H3K27me3, and H3K9me3 histone modifications in *upd*-tumor testes expressing H3-GFP in either GSC-like or CySC-like cells. In brief, *upd*-tumor testes were dissociated into single-cell suspensions, crosslinked with formaldehyde, and lysed to release chromatin. Chromatin was incubated with a ProteinA-MNase-antibody complex to direct cleavage in the cell type expressing H3-GFP. This was followed by controlled MNase digestion to release soluble fragments. Antibody-coupled magnetic beads were prepared in parallel and used to capture target chromatin, which underwent sequential low- and high-salt washes before elution. Crosslinks were reversed, DNA was purified by a spin-column cleanup, and both input and immunoprecipitated fractions quantified. Sequencing libraries were prepared using the NEBNext DNA Ultra II DNA library kit and sequenced on the Illumina NovaSeq platform.

### Replication sequencing (Repli-seq)

Repli-seq was performed on *upd*-tumor testes expressing H3-GFP in either GSC-like or CySC-like cells according to a published protocol (Marchal et al. 2018). Briefly, *upd*-tumor testes were dissected, incubated with BrdU, and dissociated to isolate nuclei. GFP-positive nuclei were sorted by flow cytometry to separate S-phase populations into four fractions (early, early-mid, late-mid, late), providing temporal resolution of replication dynamics. DNA was then extracted from ∼10,000 nuclei per fraction, fragmented by sonication, and prepared for sequencing using the NEBNext Ultra II DNA Library Prep Kit. BrdU-labeled nascent DNA was enriched by sequential immunoprecipitation with an anti-BrdU antibody followed by a secondary anti-mouse IgG, then purified after Proteinase K digestion. The recovered DNA was quantified, PCR-amplified, and pooled for sequencing on the Illumina NovaSeq platform.

### Bioinformatics

All genomic findings are identified based on alignment to FlyBase’s *Drosophila melanogaster* genome (release 6.47) (Öztürk-Çolak et al. 2024). Supplemental steps which support particular analyses are: Only analyzing uniquely-mapped ChIC-seq data (MAPQ>=20); application of epigenomic borders of the pericentromeric heterochromatin for chromosome region summaries (Filion et al. 2010); trimming rDNA-adjacent noncoding DNA from the rDNA auxiliary sequence file (to summarize or plot the rDNA repeat unit); and regressing ChIC-ChIP enrichment on the unique transposon sequence set.

#### Single-cell RNA sequencing analysis

To test the hypothesis that the GSC-like and CySC-like tumor cell populations are largely homogenous, we aligned and quantified the scRNA-seq UMI data using Cell Ranger (v7.0.0) with default parameters (Zheng et al. 2017). Our dimension reduction (UMAP) and clustering are derived from Seurat’s SCTransform integrated assay (Hafemeister and Satija 2019; Butler et al. 2018). For quantification, single-cell data is de-noised using DecontX for UMI quantification correction (Yang et al. 2020). Log normalized gene expression is computed from the DecontX corrected count for the gene, with a shifted logarithm (Ahlmann-Eltze and Huber 2023). Pseudobulk Log_10_ CPM is computed, after a compatible isoform selection, re-reapplication of DecontX, and log transformation, using a linear model (Ritchie et al. 2015). Posterior log fold change estimates further enrich the downstream gene set enrichment, compared to pseudobulk log-scaled differences, by fitting a Bayesian regression coefficient estimate (Zhu et al. 2019). Our covariates are DecontX’s contamination score, crossed with cell type.

Differentially expressed genes (DEGs) for GSC-like or CySC-like enrichment were selected by *s*-value less than 10^-4^ and log effect size (L_2_FC) at least 1.5, yielding enriched Gene Ontology (GO) terms (Thomas et al. 2022).

#### ChIC-ChIP sequencing analysis

ChIC-seq reveals nucleosome occupancy, H3K4me3 enrichment, H3K27me3 enrichment, and H3K9me3 enrichment, in the bulk GSC-like and CySC-like cell genomic DNA libraries. Our ChIC-seq analyses produces sharp nucleosome positioning estimates as regression effects (intercept of H3 abundance and L_2_FC of ChIC-ChIP treatment) (20 bp step) where the model applied is maximum-likelihood Negative Binomial regression (Ahlmann-Eltze and Huber 2021). Positioning is made to be precise by filtering paired-end fragment length and only binning the midpoint between the reads into the sliding genomic window. Monosomes are visible as local maxima in a genomic track, producing the finished ChIC-seq tracks, after estimating feature density from regression predictions: smoothing using a Gaussian kernel with sigma = 40 bp (Boyle et al. 2008). The log-fold enrichment of the IP “F-Seq” (Gaussian) track, relative to the H3 input track, reveals antibody binding to the monosome, with appropriate (20 bp) resolution to reveal the monosome midpoint location.

#### Replication sequencing analysis

Repli-seq identifies genomic loci that are enriched in a certain interval of S-phase (where the interval of S-phase is marked by molecular mass of DNA). Like our ChIC-seq regression, we bin the midpoints of Repli-seq single-end 100 bp reads or the properly paired alignment (up to 500 bp wide) into non-overlapping 1kb steps across the genome. We found that consecutive bins of fragments were statistically independent enough (in autocorrelation) to serve as a set of observations for regression. In a 3 kb window, we used the observations of 3 bins of DNA in either 6 or 8 samples to infer a Bayesian logistic regression parameter (logit timing score), characterizes by its posterior expected value. The timing parameter posterior distributions are compared to reject the null hypothesis of a static replication regime. Testing a coefficient to explain biological count data can be carried out via a likelihood ratio test (Love et al. 2014). We apply numerical integration to the full and reduced model likelihoods, summarized as a Bayes factor. For pairwise cell type comparisons, the null hypothesis is rejected using the Bayes factor in regions of width at least 20kb. Finally, for viewing and for marking early- and late-replicating chromatin, Repli-seq timing estimates can be smoothed using LOESS.

This version of the genomic tracks permits *Z*-scoring several cell types at every genomic location.

Mean *Z*-scores across the chromosomal regions allow us to describe regions as having overall static choreography or replication bias.

## DATA ACCESS

All raw and processed sequencing data generated in this study have been submitted to the NCBI Gene Expression Omnibus (GEO; https://www.ncbi.nlm.nih.gov/geo/) under the accession number GSE291929. The genomics analysis scripts are submitted as Supplemental Code and have been deposited in GitHub: https://github.com/ringw/Upd-Germline-Genomics

## ACKNOWLEDGEMENTS

We thank Dr. Winston Timp and Siqi (Alice) Chen for technical support in performing the 10x Genomics scRNA-seq, Dr. Hao Zhang (Flow Cytometry and Cell Sorting Core Facility, Johns Hopkins Bloomberg School of Public Health), for assistance with Repli-seq cell sorting, and Hanhvy Bui (John Hopkins Integrated Imaging Center) for her initial efforts in developing a cell sorting protocol. We are grateful to the Johns Hopkins Genetics Resources Core Facility for consultation and high-throughput sequencing.

We also thank several members of Dr. Keji Zhao’s group - Drs. Lixia Pan, Wai Lim Ku, Guangzhe Ge, Kairong Cui, Qingsong Tang, and Meiyuan Ji - for help in establishing the ChIC-seq protocol, sequencing submission, and data storage. Funding was provided by NIH Intramural funding (KZ); NIH R35 GM127075-01S1, American Cancer Society PF-19-131-01-DMC, and NIH MOSAIC K99GM145973 (JAU); NIH R35 GM127075, R01 HD102474, and the Howard Hughes Medical Institute (XC).

## AUTHOR CONTRIBUTIONS

Project was conceptualized by JAU and XC. Validation of the *upd-*tumor system was completed by RG and WX, while all other experimentation was performed by JAU. The bioinformatic analysis was performed by DR and JMU. This manuscript was written and edited by JAU and XC. Support for experimentation and funding for this study was provided by JAU, KZ, and XC.

