## Supplemental Materials for "Cell type- and chromosome-specific chromatin landscapes and DNA replication programs of *Drosophila* testis tumor stem cell-like cells"

**Supplemental Figure 1.** Single-cell RNA-seq of *upd*-tumor testes classifies two stem cell populations

**Supplemental Figure 2.** Comparison of cell- and stage-specific marker gene expression in GSC-like and CySC-like cells from the upd-tumor scRNA-seq dataset to that of wild-type testis snRNA-seq

**Supplemental Figure 3.** Intercellular communication *versus* chromatin-based regulation in stem cell identity

**Supplemental Figure 4.** Cell-specific profiles of H3K4me3, H3K27me3, and H3K9me3

**Supplemental Figure 5.** The replication timing profiles of GSC-like and CySC-like cells

**Supplemental Figure 6.** GSC-like cells have a distinct replication timing program from CySC-like and cell culture cells

**Supplemental Figure 7.** Differences in replication timing correlate with differences in chromatin enrichment

**Supplemental Methods**

**Supplemental Tables S1-S7,** provided as separate excel files

**Supplemental Code**, provided as a separate zipped file

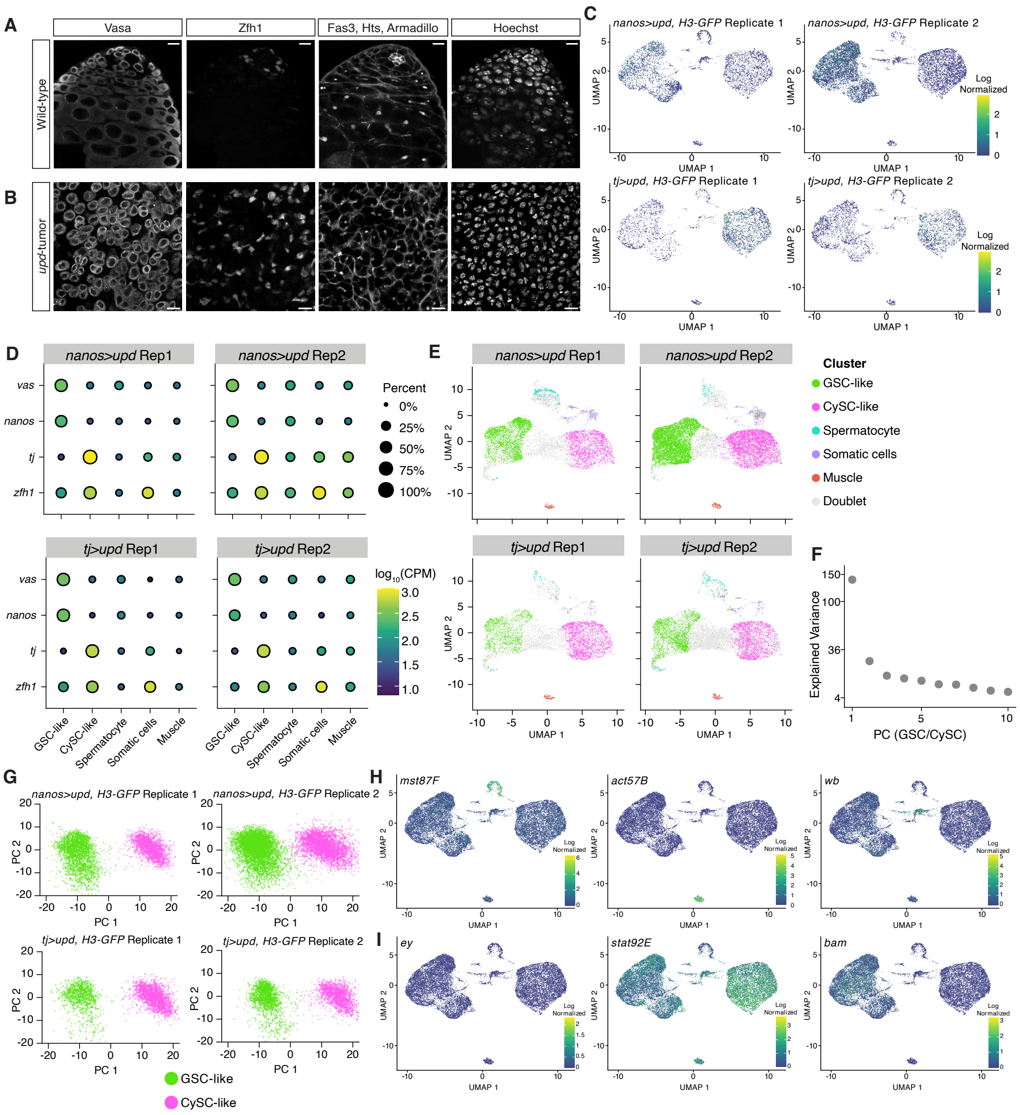


### Figure S1: Single-Cell RNA-seq of *upd*-tumor testes classifies two stem cell populations

A&B: Greyscale images of individual immunostaining panels associated with Fig. 1A&B for wild-type and *upd*-tumor testes, respectively. Vasa is a cytoplasmic protein present in germ cells. Zfh1 is a cyst stem cell transcription factor. Hts and Armadillo mark the spectrosome/fusome and intercellular connections, respectively. DNA is stained with Hoechst. Scale bar = 10 μm.

C: Uniform Manifold Approximation Projection (UMAP) graphs showing expression of the H3-GFP transgene in each scRNA-seq sample replicate after filtering out doublets. H3-GFP was expressed in either germ cells (*nanos>upd, H3-GFP*) or cyst cells (*tj>upd, H3-GFP*), as indicated.

D: The expression of cell-specific marker genes is shown as the average $\log_{10}$ (CPM) for the five cell-type clusters identified in each single-cell RNA-seq replicate. The percent of cells in a cluster with the respective transcript expressed is indicated by the size of the dot.

E: UMAP showing the detection of doublet cells in each of the scRNA-seq replicates. Integrated cells used in the scRNA-seq analysis (colored) are indicated along with the rejected (excluded) doublet cells in light grey. Grey cells failed either the scDblFinder check or the nUMI filtering that was applied prior to replicate integration. The grey cells occupy a space in the UMAP that is in between the GSC-like, CySC-like, and spermatocyte single cells.

F&G: After integrating genotypes and replicates, the majority of transcriptional variance was orthogonal to tumor genotype variable (Principal Component 1 (PC1) *vs. nanos>upd, H3-GFP/tj>upd, H3-GFP*: $R^2 = 0.06$) , as indicated by a several-fold explained variance difference vs the remaining components. PC1 effectively separates GSC-like cells from CySC-like cells along the x-axis, while not capturing directions of variance between the genotypes.

H: UMAP graphs of known marker transcripts that identify the cell-types of other clusters in the dataset.

I: Expression UMAP graphs of stage-specific transcripts showing the GSC-like and CySC-like clusters have stem-cell identity and are not differentiated. The transcript *eyeless* (*ey*), a gene required for eye development, was used as a negative control and not detected in any clusters (Halder et al. 1995).

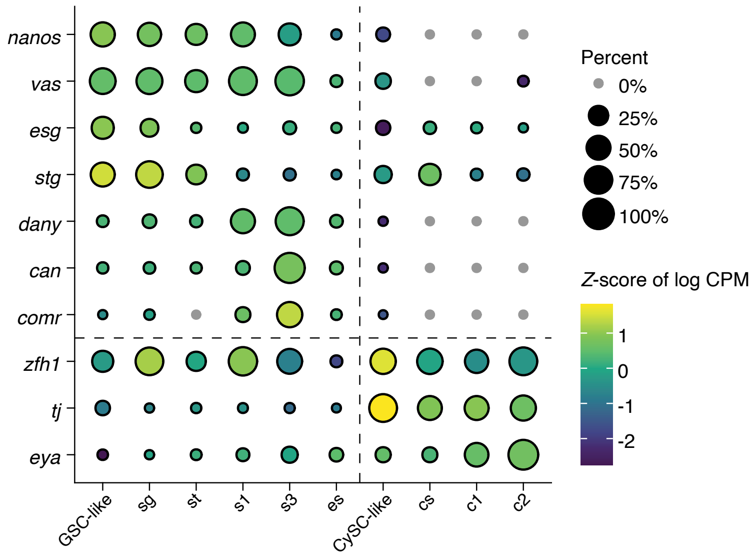


**Figure S2: Comparison of cell- and stage-specific marker gene expression in GSC-like and CySC-like cells from the *upd*-tumor scRNA-seq dataset to that of wild-type testis snRNA-seq**

Annotations for the wild-type snRNA-seq data are taken from Raz, et al (Raz et al. 2023). In addition to expressing *nanos* and *vas*, GSC-like cells from this study are transcriptionally comparable to the "sg" cluster in the wild-type dataset, which contains a mixture of GSCs and early spermatogonia due to technical constraints in isolating sufficient pure GSCs. In addition to expressing germline markers *nanos* and *vas*, GSC-like cells also express germ-stem cell transcripts *esg* and *stg* (Kiger et al. 2000; Inaba et al. 2011). By contrast, late-stage spermatocyte (st – s3) markers *distal antenna-young (dany)*, *cannonball (can)*, and *cookie monster (comr)* are absent from the GSC-like cluster, whereas elongating spermatids (es) lack the expression of all shown marker genes (Trost et al. 2016; Hiller et al. 2001; Laktionov et al. 2014). Similarly, CySC-like cells from the *upd*-tumor resemble the cyst stem cell (cs) cluster in the wild-type dataset, characterized by expression of *zfh1* and *tj*, but not the late-stage (c1 and c2) marker *eya.*

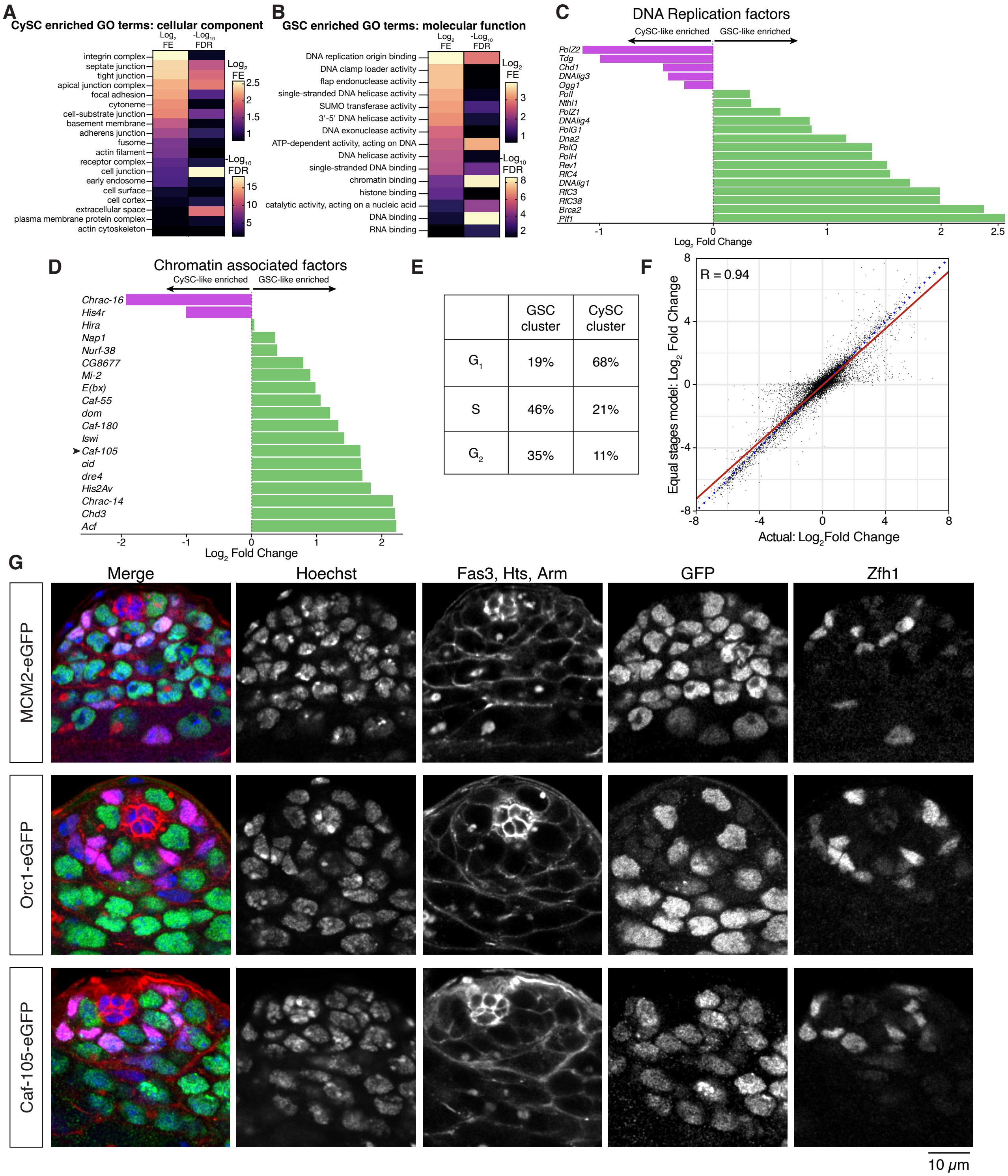


| | GSC cluster | CySC cluster |
|---|---|---|
| $G_1$ | 19% | 68% |
| S | 46% | 21% |
| $G_2$ | 35% | 11% |

**Figure S3: Intercellular communication *versus* chromatin-based regulation in stem cell identity**

A&B: Gene Ontology analysis results for cellular components (CySC-like cells) and molecular function (GSC-like cells) of significantly enriched genes identified by differential gene expression analysis. Shown are heatmaps of the $Log_2$ Fold Enrichment (FE) and negative $Log_{10}$ False Discovery Rate (FDR) results from the GO analysis.

C&D: The $Log_2$Fold Change [$Log_2FC = log_2(GSC/CySC)$] values of indicated transcripts that are involved in DNA replication or chromatin regulation.

E: Using Seurat's module score of phase-specific transcripts, the percent of GSC-like or CySC-like cells in each indicated stage of the cell cycle was calculated.

F: A correlation plot of transcript $Log_2FC$ in actual data compared with a model whereby each cell cycle stage is represented in equal parts. The high $R^2$ value indicates cell cycle stage does not affect the transcriptional profile.

G: Confocal immunostaining of non-tumor (wild-type) whole testis tissue. Shown are individual channel z-stack images of strains with the indicated endogenously tagged protein. Expression of each protein is restricted within the mitotically active spermatogonial region. Hoechst stains DNA within each nucleus. Fas3 identifies the stem cell niche. Hts and Armadillo mark the spectrosome/fusome and individual germline cysts, respectively. Zfh1 is a transcription factor expressed only in cyst stem cells.

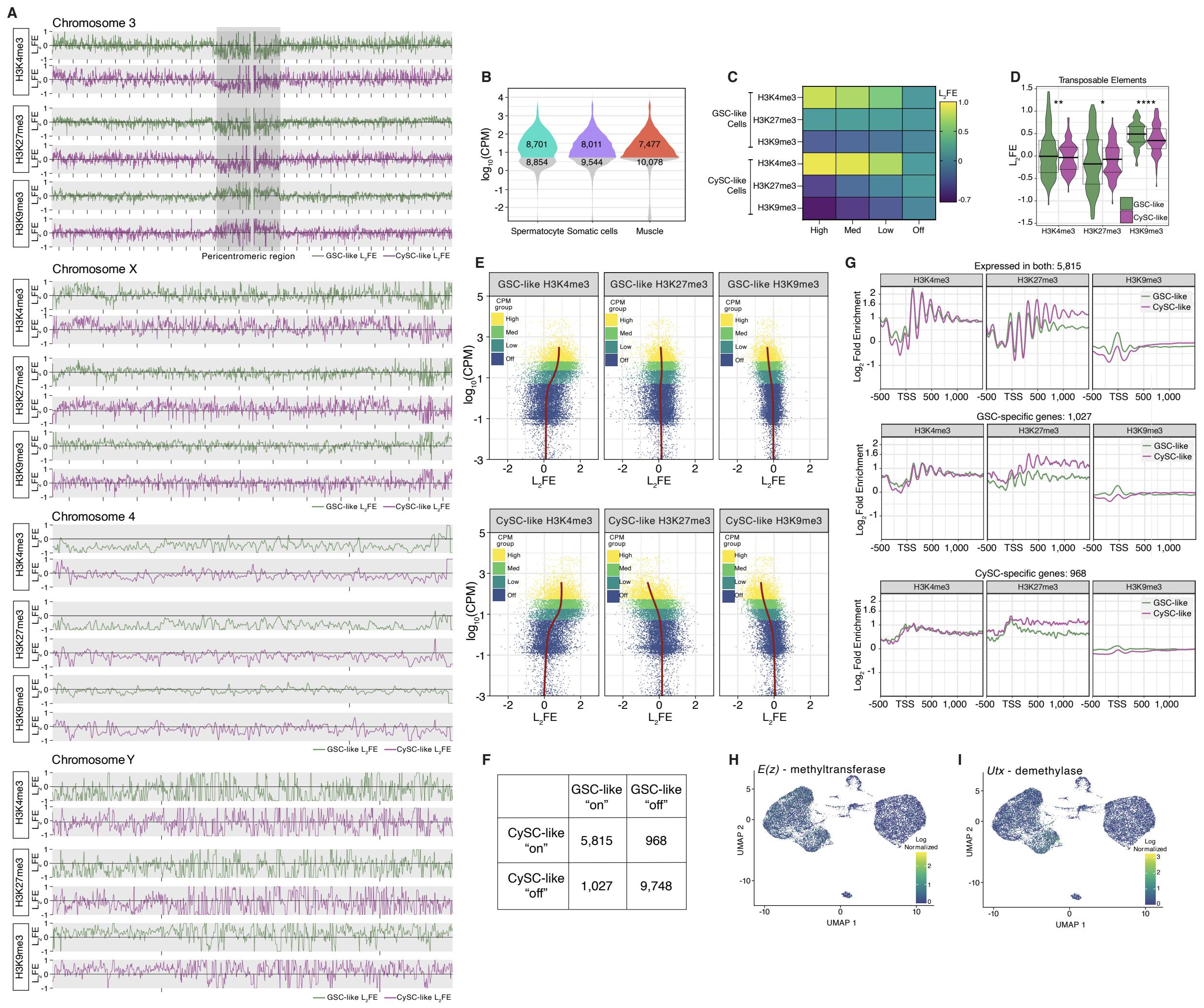


**Figure S4: Cell-specific profiles of H3K4me3, H3K27me3, and H3K9me3**

A: We quantified histone mark enrichment using a multiple regression model that includes the input log-enrichment as intercept, a ChIP $Log_2$ Fold Enrichment ($L_2FE$) coefficient, and covariates. The $L_2FE$ coefficients robustly characterize the chromatin of the *Drosophila* chromosomes including Chr3, Chr4, ChrX, and ChrY. The pericentromeric region of Chromosome 3 is indicated in grey. Enrichment tracks from GSC-like cells are in green, whereas the tracks associated with CySC-like cells are in magenta. For all chromosomes, the distance between X-axis tick marks is equivalent to 2Mb.

B: The transcriptomes of the spermatocyte, other somatic cells, and muscle cell clusters obtained from the scRNA-seq were categorized as either expressed or off using a cut-off of CPM>5 to demonstrate the similarity in abundance distributions.

C: We applied a LOESS model, regressing the histone modification $L_2FE$ at TSSs with the gene expression levels ($Log_{10}CPM$), to estimate mean $L_2FE$ values for genes categorized as 'off', 'low', 'medium', and 'high' based on their median $Log_{10}CPM$ values. Regression coefficients of H3K4me3, H3K27me3 and H3K9me3 ($L_2FE$) are summarized here.

D: Enrichment of H3K4me3, H3K27me3, and H3K9me3 at *Drosophila* transposable elements, as the distribution of $L_2FE$ coefficients, where one coefficient is computed for a window covering each transposable element reference sequence from end to end. Statistical differences in $L_2FE$ distributions between GSC-like and CySC-like transposon-wide histone mark enrichment were determined by a paired samples *t*-test (Supplemental Table 4).

E: Local regression (shown transposed) of H3K4me3, H3K27me3, and H3K9me3 $L_2FE$ coefficients at every gene's TSS, with the gene's log-quantification level as the regressor, reveals whether any variance in the quantification level (the regressor) is associated with the histone code.

F: A contingency table with the number of genes categorized as either on in both cell types or uniquely expressed. "On" genes were defined as having a CPM value greater than or equal to 5.

G: The $Log_2$ Fold Enrichment [$Log_2$(Average mark signal / H3 (input) signal)] for H3K4me3, H3K27me3, and H3K9me3 is plotted at the TSS of genes categorized as either expressed in both GSC-like and CySC-like cells or uniquely expressed in either GSC-like or CySC-like cells.

H&I: UMAP graphs show the expression levels of indicated H3K27me3 modifying enzymes.

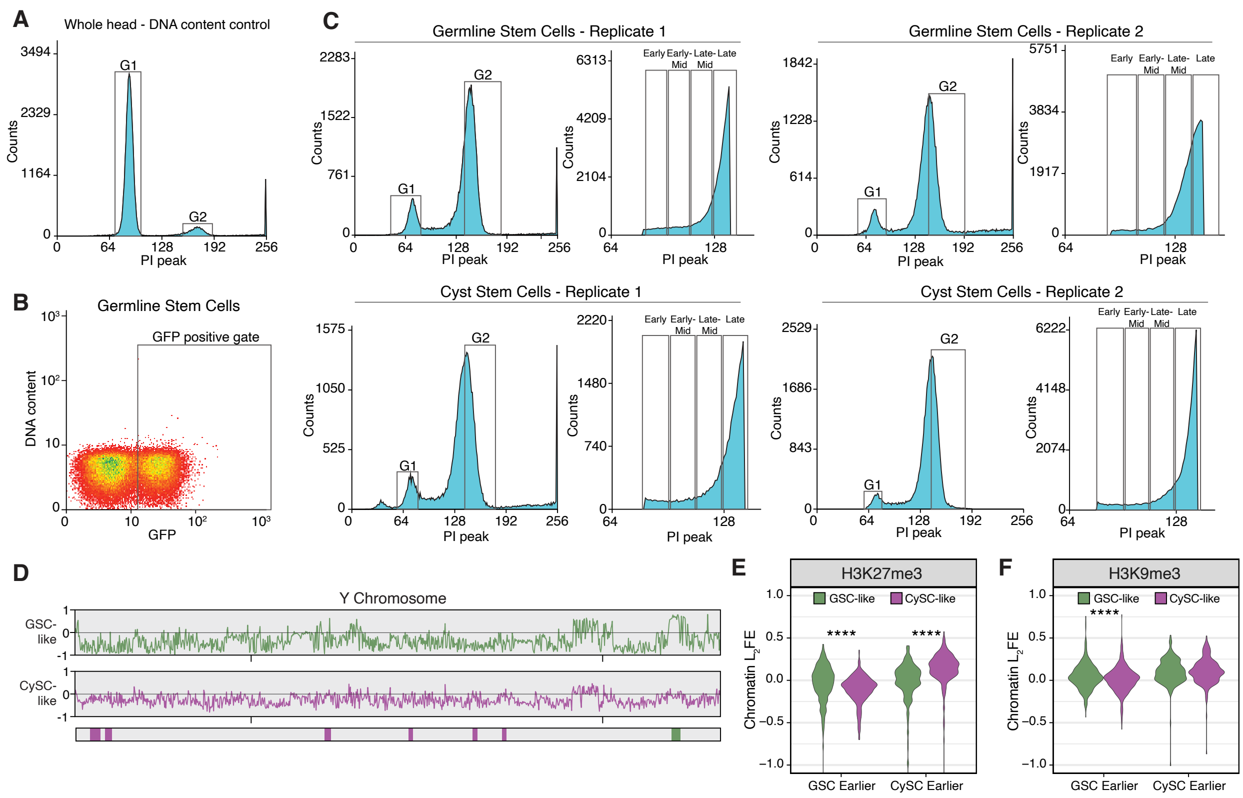


**Figure S5: The replication timing profiles of GSC-like and CySC-like cells**

A: DNA content of whole fly heads was used to determine the cell sorting gates for $G_1$ and $G_2$, as this nuclei population is predominantly in $G_1$.

B: GFP fluorescence gate used to define GFP-positive nuclei during cell sorting.

C: DNA content graphs, as measured by propidium iodide (PI), indicating the $G_1$ and $G_2$ gates. In the S-phase fraction, the four gates used to separate S-phase are shown for each replicate.

D: Replication timing for the Y Chromosome is scored as -1 (late) to 1 (early) for both GSC-like (green) and CySC-like (magenta) cells. Beneath the line shows the genomic windows where replication timing significantly differed between cell types, where the color represents the cell type with earlier replication.

E&F: The $Log_2$Fold Enrichment values for GSC-like and CySC-like earlier replicating regions are plotted for H3K27me3 and H3K9me3. Statistical differences in $L_2$FE distributions between GSC-like and CySC-like samples were determined by a paired samples *t*-test (Supplemental Table 5).

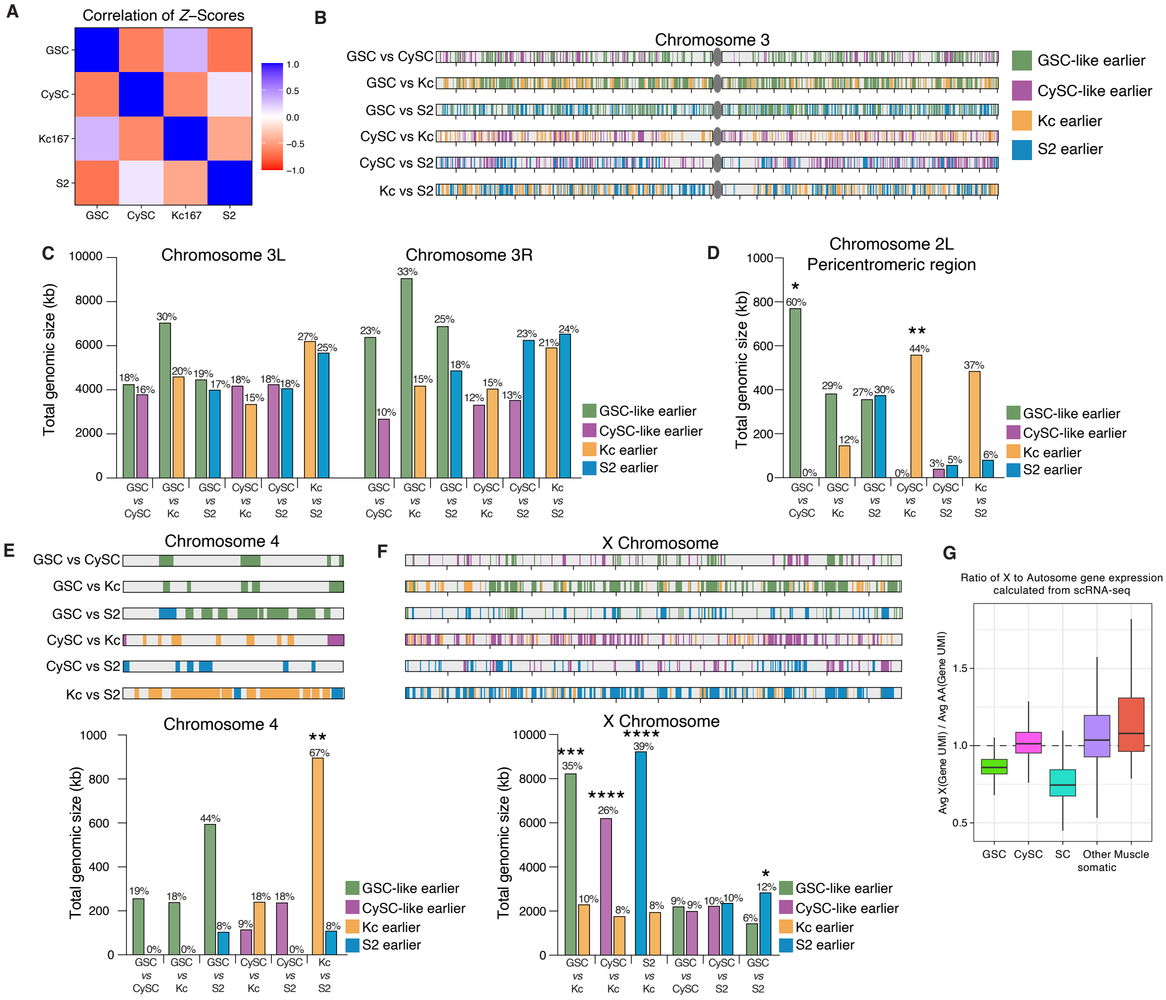


**Figure S6: GSC-like cells have a distinct replication timing program from CySC-like and cell culture cells**

A: Replication timing (RT) *Z*-scores were determined by standardizing variance at each window on the genome, then averaged across 11 chromosomal regions and scored at the ribosomal DNA (rDNA) locus, yielding cell-type profiles with a dynamic Pearson correlation. In terms of these shifts in timing of distinct chromatin locations, the GSC-like and Kc167 replication programs share significant similarities (positive correlation), as to a lesser extent do the programs of the CySC-like and S2 cells.

B: Shown is the location and identification of regions that are replicating earlier on Chromosome 3 for each indicated pairwise comparison of replication timing.

C: The total genomic size replicating earlier for each pairwise comparison is plotted for the left and right arms of Chromosome 3. There are no significant differences in the total genomic size for any comparison. Above each bar is the corresponding fraction of the chromosome that is earlier replicating, calculated as (total genomic size in bp / size of chromosome arm). Statistical significance for total genomic size was calculated using Fisher's exact test with category aggregation (Supplemental Table 6).

D: The total genomic size of replicating earlier regions within Chromosome 2L pericentromeric region are plotted for all pairwise comparisons. When compared to CySC-like cells, GSC-like cells have significantly more earlier replicating DNA. Likewise, Kc167 cells have a significantly greater total genomic size that replicates earlier than CySC-like cells (Supplemental Table 6). The corresponding percentage of the pericentromeric region that is replicating earlier is indicated above each bar.

E. The total genomic size of replicating earlier regions on Chromosome 4 are plotted for all pairwise comparisons. When compared to all other cell types, a greater amount of the $4^{th}$ chromosome replicates earlier in GSC-like cells. Another cell comparison with large differences in replication timing is Kc167 *versus* S2 cells. For all comparisons the percentage of the $4^{th}$ Chromosome that corresponds to the total genomic size with earlier replication is indicated above the bar.

F: The total genomic size, corresponding percent of the chromosome, and the location of replicating earlier regions on the X Chromosome are plotted for all pairwise comparisons. In all comparisons between a male (XY) and female (XX) cell type, the male cell has a greater amount of earlier replicating regions than the Kc167 cells.

G: Ratio of gene expression levels between X-linked and autosomal genes calculated using our scRNA-seq data for each of the five cell clusters shows no evidence for equalization of X-linked gene transcription in GSC-like cells or spermatocytes.

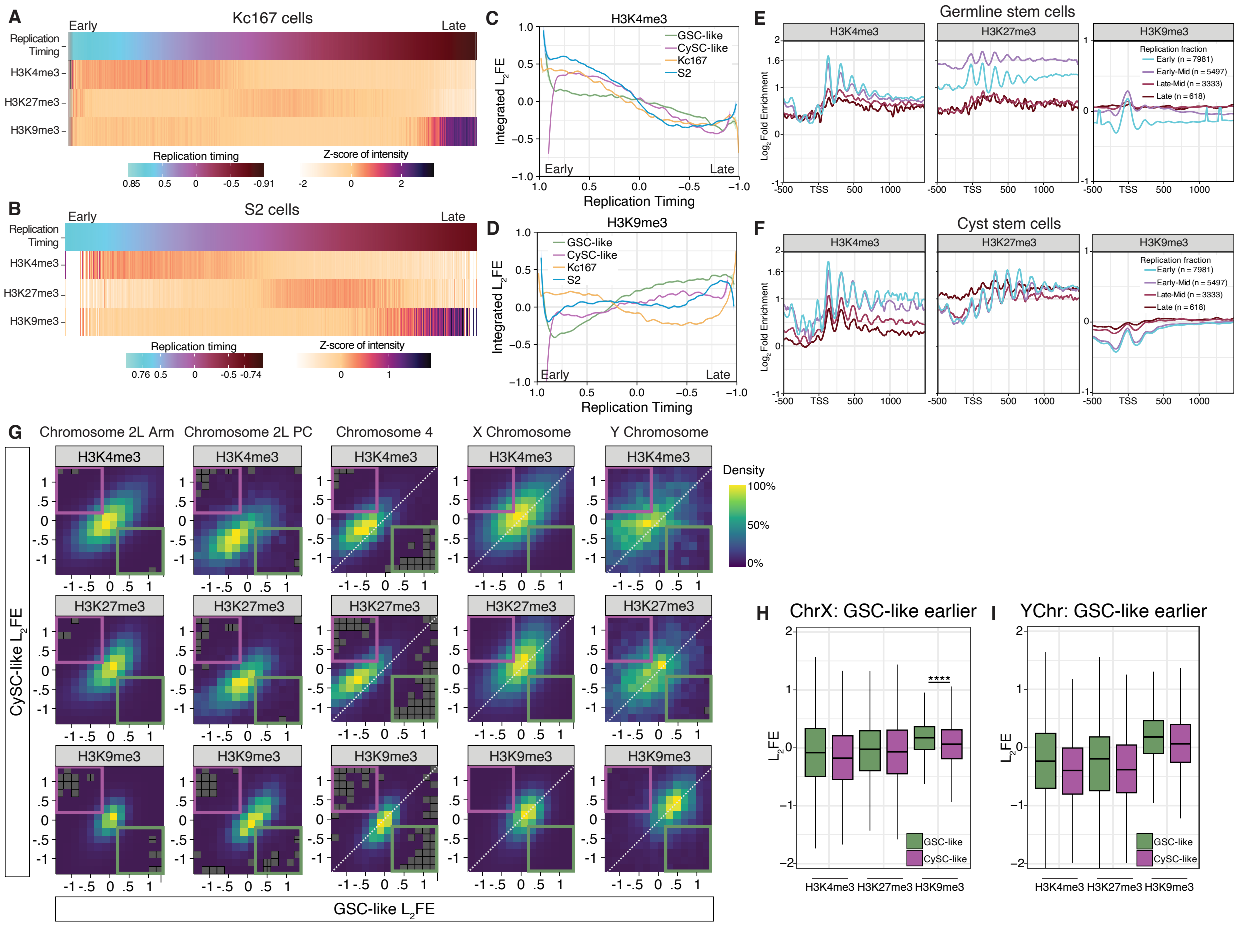


**Figure S7: Differences in replication timing correlate with differences in chromatin enrichment**

A: Histone modification enrichment from modENCODE is ordered according to replication timing for Kc167 cells. There is increased H3K4me3 during early replication and increased H3K9me3 during late. H3K27me3 is increased more during mid-S-phase.

B: Histone modification enrichment from modENCODE ordered according to replication timing for S2 cells. H3K4me3 is enriched in early replicating regions, H3K27me3 is present during mid-S-phase and H3K9me3 enriched in late replicating regions.

C: Enrichment of H3K4me3 in GSC-like, CySC-like, Kc167, and S2 cells is plotted according to replication timing, showing that for all cell types H3K4me3 has greater enrichment at regions that replicate early in S-phase.

D: Enrichment of H3K9me3 in GSC-like, CySC-like, Kc167, and S2 cells is plotted according to replication timing, showing that H3K9me3 has greater enrichment at regions that replicate late in S-phase for all cell types.
E: The $Log_2$Fold Enrichment ($L_2$FE) of H3K4me3, H3K27me3, and H3K9me3 is plotted at TSS of genes in each of the 4 categorized replication timing fractions (early, early-mid, late-mid, or late). GSC-like cells have more H3K4me3 at genes that replicate early, whereas H3K27me3 is enriched at genes that replicate during mid-S-phase.
F: The $L_2$FE of H3K4me3, H3K27me3, and H3K9me3 is plotted at TSS of genes found in each of the 4 categorized replication timing fractions (early, early-mid, late-mid, or late). CySC-like cells have more H3K4me3 at genes that replicate early, whereas H3K27me3 is at similar levels at genes that replicate during early and mid-S-phases.
G: The GSC-like and CySC-like $L_2$FE distributions (marginal distributions shown previously) have a positively correlated joint distribution. The highlighted quadrants (excluding chromatin that has a near-zero GSC-like or CySC-like $L_2$FE) reveal the GSC-enriched (green box) and CySC-enriched (magenta box) classifications. These regions have $L_2$FE values >0.2 in one cell type and <-0.2 in the other.
H&I: The $L_2$FE values for the three histone modifications in GSC-like and CySC-like cells are plotted for X and Y chromosome regions that replicate earlier in GSC-like cells. Statistical significance (*) is measured by a paired sample *t*-test (Supplemental Table 5).

#### **Supplemental Methods**

##### Fly stocks

| **Experiment Genotype** | **Parental Genotypes** | **Source** | **Experiment** |
|---|---|---|---|
| y-w- ; Vasa-eGFP / + ; | y-w- ; + ; + | Chen lab | Immunostaining, Figure 1 |
| | w- ; Vasa-eGFP ; | Kyoto Drosophila stock center | |
| w- ; Vasa-eGFP / UAS>Upd ; *nanos*>Gal4 / MKRS | w- ; Vasa-eGFP ; *nanos*>Gal4 | Chen lab | Immunostaining, Figure 1 |
| | w-; UAS>Upd ; MKRS/TM6b | Chen lab | |
| HS>FLP ; *nanos*>Gal4 / UAS>Upd ; H3V6 / + | hsFLP ; *nanos*>Gal4 ; + | Chen lab | **Germline-specific** Single-cell RNA-seq ChIC-seq Repli-seq |
| | w- ; UAS>Upd / CyO ; H3v6 | Chen lab | |
| + ; *Tj*>Gal4 / UAS>Upd ; H3v6 / + | +; Tj>Gal4 / CyO, Kr>GFP ; + | Van Doren lab | **Somatic-specific** Single-cell RNA-seq ChIC-seq Repli-seq |
| | w- ; UAS>Upd / CyO ; H3v6 | Chen lab | |
| w-; Orc1-GFP ; + | Stock | This study | Immunostaining, Figure 2 |
| w-; + ; MCM2-GFP (c-term) | Stock | This study | Immunostaining, Figure 2 |
| w-; Caf-105-GFP(n-term) ; + | Stock | This study | Immunostaining, Figure 2 |

##### Antibodies

| **Primary antibodies** | **Source** | **Dilution** | **Experiment** |
|---|---|---|---|
| Chicken anti-GFP | Abcam #13970 | 1:1000 | Immunostaining |
| Rabbit anti-Zfh1 | Ruth Lehmann | 1:2000 | Immunostaining |
| Mouse anti-Fas3 | DSHB #7G10 | 1:50 | Immunostaining |
| Mouse anti-1B1/Hts | DSHB #1B1-s | 1:50 | Immunostaining |
| Mouse anti-Armadillo | DSHB #N2 7A1 | 1:50 | Immunostaining |
| Mouse anti-Alpha-Spec | DSHB #3A9 | 1:50 | Immunostaining |
| Rabbit anti-GFP | Abcam #290 | | ChIC |
| Rabbit anti-H3K4me3 | Abcam #8580 | | ChIP |
| Rabbit anti-H3K27me3 | EMD Millipore #07-449 | | ChIP |
| Rabbit anti-H3K9me3 | Diagenode #C15410056 | | ChIP |
| Mouse anti-BrdU | BD Biosciences #555627 | | Repli-seq |
| Rabbit anti-mouse IgG | Sigma #M7023 | | Repli-seq |

| **Secondary antibodies** | **Source** | **Dilution** |
|---|---|---|
| Goat anti-chicken 488 | Abcam #150169 | 1:1000 |
| Goat anti-rabbit 568 | ThermoFisher #A11011 | 1:1000 |
| Goat anti-mouse 680 | ThermoFisher #A32729 | 1:1000 |

##### Immunostaining (whole mount method)

Testes from 1-5 day old adult male flies were dissected in pre-warmed Schneider's media (Gibco #21720-024) before fixing at room temperature in 4% formaldehyde (Cell Signaling #12606) in 1× PBS containing 0.1% Triton X-100 (PBST; Quality Biologicals #119-069-131, Fisher #BP151) for 10 minutes with rotation. Following fixation, samples were briefly rinsed twice in PBST followed by three 5-minute PBST washes with rotation. Primary antibodies were prepared at appropriate dilutions in 5% normal goat serum (NGS) (Cell Signaling #5425) in PBST and the testes incubated for three nights at 4°C.

After incubation, the primary antibody solution was removed and two brief PBST rinses performed. The testes were next washed three times in PBST for 5-minutes each with rotation. Secondary antibodies were diluted 1:1000 in 5% NGS in PBST and testes were incubated for two nights, shielded from light at 4°C with rotation. At the end of the secondary incubation, Hoechst 33342 (Invitrogen #H3570) solution was added to secondary solution at a dilution of 1:1000, and the testes rotated for 10 minutes. Afterward, the secondary antibody solution was removed and two brief PBST washes performed. This was followed with three 5-minute PBST washes with rotation. Testes were then mounted on a Superfrost Plus microscope slide (Fisher #1255015) using Vectasheild PLUS Antifade mounting media (Vector Laboratories #H-1900). Imaging was performed on a Leica Stellaris5 Confocal with 405nm and white light laser.

Immunostaining (squash method)

*Upd*-tumor testes (*nanos*>Upd) were dissected in warm Schneider's media. Approximately 5-6 individual testes were transferred to a microscope slide, and excess media removed. A 10uL drop of 1× PBS was placed over the tumors before gently rupturing each tissue. A coverslip was placed over the ruptured testes to squash and spread the tissue. Slides were then flash frozen in liquid nitrogen for 2 minutes. Following freezing, the coverslips were removed with a razor blade and the slides immediately placed in a coplin jar containing pre-chilled 95% ethanol (Fisher #BP2818). The jar was incubated at -20°C for 10 minutes. Following incubation, slides were removed from the jar. Excess ethanol was wiped from the slide and the tissue outlined with a hydrophobic pen. A 50uL drop of 1% formaldehyde in 1× PBS with 0.1% Triton X-100 (PBST) was placed on top of the tissue and covered with parafilm. The slides were fixed in a humid chamber for 3 minutes, after which the slides were rinsed quickly in PBST thrice.

Following washes, the slides were permeabilized in a PBST/DOC (1× PBS, 0.5% Triton X-100, 0.5% Sodium Deoxycholate) solution for 15 minutes at room temperature with gentle rocking. This was repeated once for a total of 2 washes. After the second wash, a PBST wash was performed for 10 minutes.

Slides were incubated with 20uL of appropriate primary antibodies diluted in 3% BSA in PBST (Cell Signaling #9998) overnight in a humid chamber at 4°C. The next day, slides were washed three times, 10 minutes each in PBST. Secondary antibodies were diluted 1:1000 in 5%NGS/PBST and incubated overnight at 4°C in a humid chamber. The following day, slides were washed three times for 5 minutes each with gentle rocking. Hoechst 33342 was added to the final wash at a concentration of 0.5ug/mL in PBST. Finally, slides were mounted in vectashield and imaged on a Leica Stellaris5 Confocal equipped with a 405nm and white light laser.

<u>Intensity quantification for GFP-tagged fusion proteins</u>

The image analysis software Imaris was used to quantify the amount of GFP signal in endogenously tagged Orc1, MCM2, and Caf-105 containing GSCs and CySCs. First, machine learning identified the nuclei of individual GSCs and CySCs as surfaces within the tissue. GSCs were determined by their adjacent proximity to the stem cell niche and presence of the spectrosome, whereas CySCs were identified using the expression of Zfh1. For each nucleus (surface), the intensity sums of both Hoechst signal and GFP were calculated using the statistics option. To normalize GFP intensity to DNA content for an individual nucleus, the intensity sum of GFP signal was divided by the intensity sum of Hoechst signal. The $\log_2$ of this value was calculated and plotted, followed by a two-tailed Mann-Whitney *U* test in GraphPad.

<u>10x Genomics single-cell RNA sequencing</u>

*Preparation of single cell solution from Upd-tumor testes for single-cell RNA sequencing*

Tumor testes from either *nanos>Upd, H3-GFP* or *tj>Upd, H3-GFP* were dissected in warm, filtered Schneider's media supplemented with 10% FBS (Thermo Fisher #16140071). After collecting ~18-20 pairs, testes were transferred to ~250uL of digestion buffer (TrypLE Express, Gibco #12605-101 with 2mg/ml collagenase, Sigma #C9407). The tissues were incubated in a 37°C waterbath, with intermittent agitation to dissociate the tissue to completion. Next, the cells were filtered first through a 40um filter (Corning #352340), followed by a 10um filter (pluriSelect, #43-10010-40). Digestion buffer was removed after pelleting the cell suspension by centrifugation at 1,200 rpm for at least 7 minutes. The cell pellet was washed twice in 500uL HBSS (Gibco #14170-112), before resuspending in 20uL HBSS. Cell count and viability was obtained at this point using Trypan blue (Gibco #15250-061), before proceeding to the 10x Genomics single cell RNA sequencing protocol.

*10x Genomics sequencing*

Single cell RNA sequencing libraries were generated following the manufacturer's guidelines for the Chromium Next GEM Single Cell 3'Kit v3.1 (#1000268). Final libraries were pooled to 4nM and sequenced on the Illumina NovaSeq sequencer, using the S1 100 cycle within the Johns Hopkins Genomics Core.

<u>Cell-specific sequential Chromatin Immunocleavage – Chromatin Immunoprecipitation (ChIC-ChIP)</u>

*Prepare magnetic beads for immunoprecipitation:*

The evening prior to dissection, immunoprecipitation (IP) beads were prepared for blocking. Per desired IP, 10uL of magnetic protein-G beads (Invitrogen, #10003D) were incubated in 200uL antibody binding buffer containing 5% BSA (10mM Tris-HCl, 1mM EDTA, pH7.5, 150mM NaCl, 0.1% Triton X-100). The beads were rotated end-over-end overnight at 4°C.

The next morning, blocked beads were prepared for immunoprecipitation by washing the beads twice with 300uL antibody binding buffer containing 0.05% BSA. To the washed beads, 1ug of ChIP antibody in 200uL 0.1% BSA in antibody binding buffer was added per IP. The beads were incubated on rotating wheel at 4°C for 6 hours while preparing tissue for dissection and chromatin extraction. Antibody information for the ChIC and ChIP experiments can be found in Supplemental Materials.

*Preparation of single cell solution from Upd-tumor testes for ChIC-ChIP-seq*

Approximately 18-20 tumor testes from either *nanos>Upd, H3-GFP* or *tj>Upd, H3-GFP* were dissected and dissociated as outlined for single cell RNA sequencing. However, instead of resuspending the cell pellet in HBSS following removal of digestion buffer, cells to be used for ChIC were resuspended in 400uL of Schneider's media with 10% FBS. To this, 16% formaldehyde (Thermo #28906) was added at 1/15 the total volume to obtain a 1% final concentration. The cells were fixed by end-over-end rotation for 5 minutes at room temperature. The fixation reaction was quenched by the addition of 1/10 volume of 1.25M glycine and additional incubation for 5 minutes. Following this, the cells were pelleted by centrifugation for 7 minutes at 1,320 rpm at 4°C. Afterward, the fix solution was removed and the cell pellet washed with 500uL cold, sterile 1× PBS. The 1× PBS wash was repeated once, after which all liquid was removed. The cells were resuspended in 20uL 1× PBS to obtain a cell count.

*Preparation of ProteinA-MNase-Antibody complex and chromatin*

For a single IP, 4uL antibody binding buffer, 1uL anti-GFP antibody, and 3uL ProteinA (PA)-MNase were prepared. The cocktail was incubated on ice for 30 minutes. Meanwhile, chromatin from formaldehyde fixed cells was prepared for binding by the PA-MNase-Ab complex by treating cells with 500uL RIPA buffer (10mM Tris-HCl, 1mM EDTA, 150mM NaCl, 0.2% SDS, 0.1% Sodium Deoxycholate, 1% Triton X-100) for 10 minutes at room temperature. Following incubation, cells were pelleted by a 5-minute centrifugation at 3,000 rpm and the RIPA was removed. The cell pellet was washed once with 500uL antibody binding buffer, before resuspending in 100uL antibody binding buffer.

*Perform PA-MNase-Ab incubation, cell washes, and prepare antibody-coupled beads*

The 100uL of resuspended fixed cells were added to the microcentrifuge tube containing 8uL of prepared PA-MNase-Ab complex. After gentle mixing, the tube was incubated on ice for 60 minutes.

Following the incubation, excess PA-MNase-Ab was removed through a series of washes. First, antibody binding buffer was removed after spinning cells at 3,000 rpm for 5 minutes at 4°C. The cell pellet was then washed with 500uL high salt wash (10mM Tris, 1mM EDTA, 400mM NaCl, 1% Triton X-100), incubating for 1 minute at room temperature before spinning. The high salt wash was repeated for a total of three washes. Next, the cell pellet was washed with 200uL rinsing buffer (10mM Tris pH 7.5, 10mM NaCl, 0.1% Triton X-100), immediately spinning down for 5 minutes. After removing the rinsing buffer, the pellet was resuspended in 40uL RSB (20mM Tris pH 7.5, 10mM NaCl, 2mM $CaCl_2$, 0.1% Triton X-100).

The blocked, antibody-coupled beads were washed simultaneously with the cell pellets. First, beads were collected and unbound antibody removed. The beads were then washed a total of three times with 500uL antibody binding buffer, rotating end-over-end for 5 minutes at 4°C. After the last wash, beads were resuspended in 10uL of rinsing buffer per IP.

*MNase digestion:*

To perform the MNase digestion, the 40uL of cells in RSB were placed in a 37°C water bath for 3 minutes. Afterward, the reaction was stopped by adding 80uL of stop buffer (20mM Tris pH8, 10mM EGTA, 20mM NaCl, 0.2% SDS). Following digestion, the insoluble chromatin was spun down at 13,000 rpm for 10 minutes at 4°C. The soluble chromatin (~120uL) was moved to a new tube, where protease inhibitor (Cell signaling #5872) was added to final concentration of 1×. Next, 5% of the soluble chromatin was reserved as the input fraction.

*Chromatin IP, Bead preparation, and washing:*

For each IP, 10uL of antibody-bound beads was added and the mixture incubated overnight at 4°C with end-over-end rotation.

The next day, the IP solution was cleared on a magnetic rack and the supernatant removed. The protein bound beads were then washed three times by adding 500uL of low salt IP wash (20mM Tris-HCl pH8, 150mM NaCl, 0.1% SDS, 2mM EDTA, 1% Triton X-100) to the beads and incubating at 4°C with end-over-end rotation for 5 minutes. After the three low salt IP washes, the beads were washed once with a high salt IP wash (20mM Tris-HCl pH8, 500mM NaCl, 0.1% SDS, 2mM EDTA, 1% Triton X-100). Following the washes, DNA was eluted from the beads by adding 150uL of ChIP elution buffer (1% SDS, 0.1M Sodium Bicarbonate) to each IP and incubating for 30 minutes in a 65°C thermomixer with vortexing at 1,200 rpm. After separating the beads on a magnetic rack, eluted DNA was transferred to a clean DNA LoBind tube. At this point, 150uL ChIP elution buffer was added to the 5% input samples. For all input and IP samples, fixed crosslinks were reversed by adding 5M NaCl (final 187mM NaCl) and 40ug Proteinase K (NEB #P8107S) and incubating overnight at 65°C.

*Purify immunoprecipitated DNA and Input samples:*

The DNA from Input and IP samples was purified using a Qiagen MinElute Reaction Cleanup Kit (Qiagen #28204) with the following modification. First, 5 volumes of ERC (750uL) were added to each sample, mixed well, and transferred to a Qiagen MinElute spin column. The sample was centrifuged at 13,000 rpm for 1 min, and flow through discarded. Next, the column was washed with 700uL PE and centrifuged once again. After transferring the column to a new DNA LoBind tube, 10uL 10mM Tris, pH8.0 was added to the spin column. Following a 1 minute room temperature incubation, the sample was eluted by spinning at 13,000 rpm for 1 minute. Eluted DNA was quantified using the Qubit dsDNA high sensitivity kit (Thermo #Q33231) before proceeding to library preparation using the instructions included in NEB Next Ultra II DNA Library Prep Kit for Ilumina (NEB #7645). Finalized libraries were sequenced on a NovaSeq sequencer at the National Institutes of Health.

<u>Replication-sequencing (Repli-seq)</u>

*Nuclei isolation, sorting, and DNA purification*

Tumor testes from either *nanos>Upd, H3-GFP* or *tj>Upd, H3-GFP* were dissected in warm, filtered Schneider's media, to obtain 100-200 total pairs. For downstream sample preparation, only tumors shaped like a large ball were collected. The testes were then incubated with rotation for 15 minutes in Schneider's media containing 100uM BrdU.

After the BrdU incubation, the Schneiders' media was removed, and the testes were washed once with 1× PBS. Nuclei isolation was performed following a published protocol, with experimental considerations (McLaughlin et al. 2022). The dissected testes were first resuspended in 1mL homogenization buffer (250mM Sucrose, 10mM 1M Tris, pH 8.0, 25mM KCl, 5mM $MgCl_2$, 0.1% Triton X-100, and freshly added 1× protease inhibitors (Cell signaling #5872) and 0.1mM DTT – omit RNase Inhibitors) and transferred to a 1mL dounce homogenizer. The tissues were dissociated with 20 strokes of the loose pestle (A), followed by 40 strokes of the tight pestle (B) before pipetting gently ~20 times with wide bore tips. The nuclei were next filtered through a 40um cell strainer, (Corning #352340), followed by a 10um filter (pluriSelect, #43-10010-40) into a fresh tube. The nuclei were then spun down at 1,000×g for 10 minutes at 4°C to remove the homogenization buffer. Following pelleting, nuclei were resuspended in 400uL resuspension buffer (1× PBS, 0.5%BSA), containing final concentrations of 250ug/mL RNase A (NEB #T3018L) and 50 ug/mL propidium iodide (Sigma #P4864). The nuclei were incubated on ice, in the dark for at least 30 minutes before proceeding to the cell sorter.

Nuclei were sorted on a MoFlo XDP cell sorter (Beckman Coulter) by gating for GFP+ cells and dividing S-phase cells into 4 equal fractions corresponding to the Early, Early-Mid, Late-Mid and Late S-phase timepoints. Total number and volume of sorted nuclei was noted.

Following the sort, components of SDS-PK were added individually to keep volume low, allowing for accurate pipetting of 10,000-20,000 nuclei per fraction (SDS-PK: 50mM Tris-HCl pH8, 10mM EDTA, 1M NaCl, 0.5% SDS, 0.2mg/mL Proteinase K). Samples were incubated at 56°C for 2 hours. Afterward, nuclei were divided into separate tubes containing ~10,000 nuclei each. The DNA from ~10,000 nuclei was then purified following the protocol for cell suspensions and Proteinase K digested samples from the Zymo Quick DNA MicroPrep kit (Zymo #D3021). Purified DNA was stored at -20°C.

*Fragmentation and library construction:*

The preparation of samples for Repli-seq mostly followed published protocols, with some modifications (Marchal et al. 2018). For each of the four S-phase fractions, two tubes of 10,000 nuclei were processed as technical replicates for Repli-seq. After thawing, the volume of DNA was brought to 100uL with UltraPure water (UPW, Invitrogen, #10977-015) and samples transferred to Diagenode 0.5mL microtubes for the Bioruptor Plus (Diagenode #C30010013). The DNA was then sonicated in a cold waterbath for 1 hour on low intensity (30sec on, 90 sec off), checking every 15 minutes for ice level. The sheared DNA was then concentrated to 15uL of UPW using the DNA Clean & Concentrator-5 kit (Zymo Research #D4003) following the instructions for DNA fragments.

The concentrated DNA was then brought to a volume of 50uL with UPW prior to starting library construction using the NEBNext Ultra II DNA Library Prep Kit for Illumina (NEB #7645). The NEBNext End Prep (step 1) and Adapter Ligation (step 2), were performed following the kit instructions. After incubation with the USER enzyme, the DNA was purified again using the DNA Clean and Concentrator-5 kit following the instructions for DNA fragments and eluting in 50uL UPW.

*Immunoprecipitation of BrdU containing DNA:*

To the 50uL of clean adapter ligated DNA, 450uL of TE (10mM Tris pH8, 1mM EDTA) was added. The DNA was then denatured by heating at 95°C for 5 minutes, followed by cooling on ice for 2 minutes. The denatured DNA was then transferred to a tube prepared with 60uL 10× IP buffer (100mM Sodium Phosphate, pH 7, 1.4M NaCl, 0.5% Triton X-100). To this, 40uL of 12.5ug/mL anti-BrdU antibody (BD Biosciences #555627) was added and the sample rocked at room temperature for 20 minutes. Next, 20ug of rabbit anti-mouse IgG antibody (Sigma #M7023) was added and the sample rocked again at room temperature for 20 minutes. Following this, tubes were centrifuged at 16,000×g for 5 minutes at 4°C to pellet the BrdU containing DNA. After removing the supernatant, the DNA pellet was washed with 750uL pre-chilled 1× IP buffer (10mM Sodium Phosphate, pH 7, 140mM NaCl, 0.05%

Triton X-100). Centrifugation was repeated and the supernatant removed. The DNA pellet was then resuspended in 200uL digestion buffer containing 0.25mg/mL Proteinase K (50mM Tris-HCl, 10mM EDTA, 0.5% SDS) and incubated overnight at 37°C in an air incubator. The next day, 1.25uL of 20mg/mL Proteinase K was added to each tube before incubating for 60 minutes at 56°C. Next, the immunoprecipitated DNA was purified using the DNA Clean & Concentrator-5 kit following the instructions for DNA fragments and eluting in 16uL UPW.

*Measuring BrdU-IP efficiency and library amplification:*

Using a serial dilution of known DNA concentrations, the amount of BrdU immunoprecipitated DNA was quantified using a standard qPCR protocol with included melt curve. The calculated total amount of adapter ligated DNA was then used to determine the number of PCR cycles required for library amplification. PCR enrichment of DNA was performed following the instructions for the NEBNext Ultra II DNA Library Prep Kit for Illumina. The final libraries were pooled to 4nM and sequenced on a NovaSeq X Plus.

#### **Bioinformatics**

##### Reference sequences

Our genomic sources for all analyses are: the FlyBase dm6 chromosomes (release 6.47) (Öztürk-Çolak et al. 2024); the sequence for H3-GFP added to the genome of the flies of both genotypes; an interval of FlyBase Chromosome 2L which serves as our reference for the histone unit tandem repeat; and/or the FlyBase transposon sequence set (*Drosophila* genus). Usage of each of these genomic resources is detailed below for each genomic result. Repetitive sequences (the transposon sequence set, as well as the histone unit tandem repeat) were analyzed in the bulk sequencing after applying RepeatMasker (version 4.1.6) to the dm6 genome (Smit et al.).

The dm6 arms of Chromosomes 2 and 3 were annotated as having mostly euchromatic left and right chromosome arms, as well as having a pericentromere (indicated with “C” or “Cen” suffix). Our pericentromeric region is extended to the Crick end of the dm6 2L and 3L, and to the Watson end of the dm6 2R and 3R (imputing pericentromeric classification where the coverage of the epigenomic pericentromeric model was sparse). We chose “classic heterochromatin” boundaries that generally fit well to the observed chromatin landscape of the GSC-like and CySC-like cells, and which furthermore were coordinates estimated from another cell line whose bulk genomics we characterized, the Kc167 cell line (Filion et al. 2010). Where these regions are plotted, the mostly euchromatic chromosome arms not in this region have their values given as the 2L, 2R, 3L, and 3R values.

The dm6 rDNA assembly includes rDNA tandem repeat units as well as intergenic DNA (Öztürk-Çolak et al. 2024). For summarizing the bulk genomics of the rDNA loci, we queried the rDNA auxiliary sequence file for the coordinates of the rRNA-coding genes, then took the start and end coordinates covering this gene-rich assembly and rounded the width of this rDNA window to one significant figure (rDNA: 36901-76900). The rDNA assembly includes several rDNA tandem units. We plot a genomic result on the rRNA-coding rDNA tandem unit with the FlyBase annotation ID CR45847, also plotting our result for 2 kb of intergenic DNA before and after the rDNA gene.

Single-cell RNA sequencing analysis

Single-cell RNA sequencing results for the *Upd* tumor were aligned and quantified using Cell Ranger version 7.0.0 (Zheng et al. 2017). For this transcriptome analysis, we prepared a genome reference concatenating the FlyBase chromosomes, release 6.47, to the H3-GFP reference sequence. Each single-cell RNA-seq sample (n = 4) was quantified with the “include-introns” option enabled. All sequencing data was analyzed in R (version 4.3.3), reading and analyzing 10x Genomics feature matrices using Seurat (version 5.0.3) (Satija et al. 2015). The filtered cell barcode matrix was normalized using SCTransform (version 0.4.1) (Hafemeister and Satija 2019). The SCTransform workflow computes residuals from a Negative Binomial high-throughput sequencing regression model, and this normalization result is then passed to a nonlinear model (Seurat integration) to remove much of the within-batch effect (Butler et al. 2018).

We noticed that in each scRNA-seq sample, there were a small percent (abundance varying between samples) of droplets with a very high number of mapped UMIs (nUMI), and that cells which were high in nUMI clustered together. Indeed, in each sample, problematic cells (in terms of outlier nUMI or in excessive UMIs co-occurring for genes *Nanos* and *Traffic Jam*) clustered into 1-2 clusters (by Seurat's clustering on SCTransform followed by PCA). In each sample, one cluster's median nUMI was more than double a typical nUMI value (the median of median nUMIs), and the cluster's cells expressed both *Nanos* (GSC-like) and *Traffic Jam* (CySC-like). Therefore, we removed the cluster after calling it a doublet or multiplet cluster. In one sample, *nanos*-Upd_H3-GFP_Rep2, we removed an additional doublet cluster which was not an outlier in nUMI but was classified as doublet by the "find doublet clusters" workflow of scDblFinder (version 1.16.0). To retain most cells in "good" clusters, while clarifying cell identity by only analyzing cells that are more similar in sequencing depth, we next applied the following filters: nUMI between 2200 and 7500, and nFeature (number of genes having a UMI mapped to the gene) between 500 and 2500.

Next, the samples (n = 4) were integrated using Seurat's IntegrateData procedure, producing an "integrated assay" where the SCTransform-normalized gene expression values are warped to optimally align the samples spatially with one another before merging the data. To find the cell identities, we applied PCA (with 7 principal components, given the reduced complexity of the Unpaired tumor system) and Seurat's clustering (with resolution of 0.1, a unitless clustering parameter that varies with number of clusters). This produced 6 clusters, of which 2 formed one large mass, expressed the GSC markers *nanos* and *vas*, and were merged to form the GSC identity. Each other cluster was treated as a distinct identity: CySC, unknown germline, unknown somatic, and muscle.

For quantification and visualization of gene expression, we applied the DecontX library (version 1.0.0) to each filtered feature matrix (n = 4) (Yang et al. 2020). The decontamination procedure updated genes such as *roX1*, where the lowest % expression (in the GSC-like cluster) was reduced from 26% to <1%. From this effect, we concluded that the UMIs in each cell were affected by ambient RNA in suspension, leading to a small perturbance of the transcriptome profile of the cell. We then applied

Seurat's NormalizeData procedure, as its Log-Normalized values have a simpler explanation (pseudo-log transform of counts per 10,000) compared to SCTransform normalization. Log-Normalized decontaminated counts are presented as the gene expression level for single cells.

We created a naive gene quantification table from the DecontX counts (CPM - sum of DecontX count values for the gene, per million DecontX count values) and found that there was a gene CDS length-related effect on gene abundance. The correlation between log-grand-total-CPM (computed on all cells) and log-CDS-length was 0.24. To ameliorate this effect, we produced a feature matrix from Cell Ranger's TX tag (listing each compatible transcript id - isoform - for each UMI) and cell barcode, while also filtering the BAM file to only view exon-aligned UMIs. We found that log(TX count) varied with log(mRNA length) (as a fixed effect in a mixed linear model, considering the factor of genes as a random effect). To pick the most likely dominant isoform, we chose the TX id which maximizes log(TX count / mRNA length) for each gene. These suggested isoform UMIs were placed in a feature matrix, filtering the original UMIs found in the Cell Ranger feature matrix, and are then cleaned up using DecontX. Next, gene-cluster CPM estimates in each scRNA-seq sample (n = 4) were used (with log-transform) to produce final CPM estimates with limma's linear model (version 3.56.2) (Ritchie et al. 2015). The correlation between filtered-by-isoform UMI log-grand-total-CPM and log-CDS-length is reduced to 0.13.

Although DecontX does decontaminate cluster-gene quantification estimates, these values are transformed from discrete (UMI counts) to continuous, and too much noise may have been removed for further regression (fold-changes for the gene can be implausibly large). For differentially expressed gene (DEG) analysis, we applied an intermediate DecontX result, the % contamination estimate for each cell, along with the filtered UMIs, for regression on biological count data. The R formula ~ *(cluster + batch) * decontXcontam* allows the contrast between clusters for the gene (log-fold change) to grow if the gene expression pattern within each batch can be explained by the column of cell contamination percentages. The additional input to regression is the size factor estimate for the cell, calculated using the deconvolution method in scuttle (version 1.12.0). glmGamPoi (version 1.14.3) fits a maximum likelihood regression model to our counts matrix (Ahlmann-Eltze and Huber 2021). Maximum likelihood regression coefficients can still contain unusably large log-fold-change values (if a gene is barely expressed). Finally, apeglm (version 1.24.0) adds an Empirical Bayes prior to regression for log-fold change shrinkage, producing our robust (maximum a posteriori) log-fold change estimates (Zhu et al. 2019). We selected the apeglm model to robustly estimate log-fold change, although it is different from our germline quantification / somatic quantification workflow. The Bayesian null hypothesis covers the region of posterior probability where the effect size ($L_2FC$) is zero, or opposite to the estimated effect's sign (local false sign rate). We can reject the null hypothesis in a two-tailed manner (the estimated effect may be negative or positive). Posterior probabilities are transformed so that we can control the false discovery rate (an *s*-value). We specifically examined genes where we rejected the null hypothesis with either a positive (GSC-enriched) or negative (CySC-enriched) $L_2FC$ sign and with $s$-value $< 10^{-4}$, as well as where the absolute effect size is at least 1.5 (visualized as a volcano plot).

Our CPM values are highly clear and interpretable, with genes that should not be expressed in a cluster (e.g. *ago3* in somatic) having CPM < 5. Therefore, for each cluster, we classify genes with CPM < 5 as off. The "on" genes are split into low, medium, and high classifications each containing 1/3 of the genes.

Finally, to best align the interpretation of principal components analysis with the experiment goals (purifying either H3-GFP-expressing GSC-like cells or H3-GFP-expressing CySC-like cells), we report a PCA model of these two large clusters alone. This purifies the integrated data (both genotypes) for both cell types of interest *in silico*. The sources of variance in two-category data appearing in principal components are within-GSC-like heterogeneity, within-CySC-like heterogeneity, and the cell type enrichment effect (a law of total variance for two groups). The alignment of the two categories to the PC axes, after removing other sources of variance (from other categories of cells), reveals whether a PC can be interpreted as a cell type difference enrichment effect (sorting the two categories of cells on one axis), or shows a spread of the cells without separating the two categories of cells. Then, an elbow plot compares the PC that sorts the two cell types on the x-axis to all other PCs and shows the success of the protocol in yielding a low-complexity system, where the between-cell-type differences explain a great deal of the complexity of the tissue.

Transcriptome Cross-Comparison Analysis

The published dataset used for the transcriptome analysis is:

1-day testis snRNA-seq, 10x Genomics technology: E-MTAB-10519

We compared the *Upd* tumor transcriptome cell types to a deeply sequenced single-nucleus transcriptome of the corresponding wild-type tissue: the dissection of 1-day wild-type testes for the Fly Cell Atlas using 10x Genomics technology (Raz et al. 2023; Li et al. 2022). Wild-type (WT) nuclei (10x Genomics technology-specific) and their filtered UMI counts were accessed using the Fly Cell Atlas's processed data in H5AD format. Like the *Upd* tumor experiment, the WT single nuclei were batched by their 10x Genomics well (3 biological replicates), and then each WT biological replicate's filtered single-nuclei matrix was further filtered using the DecontX library. Each decontaminated single-nucleus experiment was pseudobulked by summing filtered UMI count estimates, log-transformed and scaled to $\log_{10}$(CPM), and we estimated each cell type's $\log_{10}$(CPM) using the limma linear model. A dot plot shows gene expression within cell types detected *in silico* in the two experiments. The plotted point color corresponds to the $\log_{10}$(CPM) (for one gene, *Z*-scored so that the individual cell type readings have a mean of 0 and a standard deviation of 1). The plotted point's radius corresponds to decontaminated percent expression rate, ranging from 0% of cells in the cell type having a UMI aligned to this gene, to 100% of the cells.

Cell Cycle-Transcriptome Analysis

Cells were classified by cell cycle (into $G_1$, S, and $G_2$M levels) using Seurat's cell cycle scores of Log Normalized gene expression (from the UMI feature matrix produced by Cell Ranger). S-phase and $G_2$M-phase scores are features combining gene sets of phase-specific transcripts, and we constructed these scores using the Tinyatlas of phase-specific transcripts (Web citation: Kirchner, R. & Barrera, V.

Drosophila_melanogaster. *GitHub*. Retrieved from https://github.com/hbc/tinyatlas/raw/add6f25/cell_cycle/Drosophila_melanogaster.csv (2024)). Crossing the major cell clusters (GSC and CySC) with classified phases ($G_1$, S, and $G_2M$) to select subsets of cells, the between-cell-type effects ($L_2FC$ in generalized linear model explaining groups) were greater than the between-phase effects in cell phase-related genes of interest such as *Orc1*. As the scRNA-seq and ChIC-seq experiments at present are asynchronous experiments, we sought to keep the scRNA-seq quantification asynchronous *in silico*. Instead, we produced an alternative version of our final and asynchronous apeglm generalized linear model using subsetting. For each 10x Genomics sample and for each cluster, we selected the phase ($G_1$, S, or $G_2M$) with the smallest number of cells, and then uniformly down-sampled cells in the other factor levels so that the *in silico* detected cell phases have abundances of 33/33/33%. Then, GLM inference proceeded on the selected cells (most cells were still retained in this alternate model). The 33/33/33% model has a general trend, on the diagonal, of not updating most transcripts' log-fold-changes very much. Although cell-phase marker genes do exist, and some transcripts' absolute log-fold-changes did shrink to less than 1, the selection based on cell phase did not significantly alter the differential expression results (R = 0.94).

#### ChIC-ChIP-seq Analysis

ChIC-ChIP sequencing results were aligned once to the dm6 genome, and once to the unique sequences (histone tandem repeat unit and transposon sequence set) reference, using Bowtie 2 (version 2.4.5) (Langmead and Salzberg 2012). The aligned reads were filtered for proper pair fragments and for a minimum MAPQ of 20 (e.g. uniquely mapping). Next, the *Drosophila* genus transposons were filtered using H3 quantification (described below). Detected transposons (n = 119) had an H3 FPKM quantification at least 0.5 times the median genome-wide H3 quantification score and were summarized in the transposon analyses.

To analyze ChIC-seq samples, we quantified fragments using a variety of sliding windows. Our method requires paired-end sequencing of digested chromatin. For all reads, we performed QC by establishing requirements for MAPQ >= 20 and SAM flag marking a proper pair (mates aligned in concordant directions). Next, an alignment of first mate and second mate genomic coordinates was counted as a single fragment, using the samtools markdup command. As our H3 input fragments had a mode length of 156 bp (GSC) or 159 bp (CySC), the fragments (after filtering to length between 100 and 200 bp) generally represent a single nucleosome, and nucleosome position can be estimated using the single base pair centered between the two mates. These base pairs are binned using sliding windows across the genome. In our ChIC-seq regression, we estimate the intercept (H3 input log-enrichment), control vs treatment (log-fold change in the mark antibody ChIP treatment), and batch effect: depending on the biological library that the input and IP come from, there is a random batch effect added to the intercept (batch effect coefficients are constrained to sum to zero). This design was chosen as each experiment has replicates, and the replicate input and ChIP are paired. The variable being regressed is the count of fragments tested against a fixed-sized sliding window on the reference.

We fitted a GLM for peak calling and quantifying $\log_2$-fold enrichment of the histone modification (using a sliding window with 500 bp width and 100 bp step). Where the monosomes are plotted, a window of 40 bp width and 20 bp step was applied instead (to identify monosome positioning). The transposon sequence set did not use a sliding window but used a single window covering each entire sequence. The fragment counts matrix entries were given offsets equal to each sample's median entry on the 2L, 2R, 3L, 3R, and 4 sequences (the autosomes), times the ratio of the observation's width in bp to the median width (which is our selected 500 bp or 40 bp window). This establishes a reference autosome-normalized level of 0 for the Negative Binomial regression with log link (the offsets matrix is log-transformed). The regression fits a fixed slope (H3 vs antibody IP L2FE) and uses sum contrasts for the factor of biological replicate, for IPs that are paired to an MNase-seq H3 input (the intercept, representing H3 quantification). Thus, the intercept is mean-centered with respect to the replicates, because the batch effects being fitted are constrained to sum to zero. The fixed batch effects that sum to zero may be considered a simplification of a generalized linear mixed model, where the batches of biological library (input and IP) are assumed to be drawn from a broader population. The generalized linear mixed model could help refine the intercept (input) quantification, because the offsets for each biological library would be shrunken to follow a normal prior (random effect).

Regression yields response predictions (expected value of H3 abundance and IP abundance, as a multiple of median autosome abundance) for all downstream analyses. However, these fitted profiles could be sparse when analyzing the 40 bp window of fragment midpoints which we expect to reveal monosome positioning. F-Seq converts from fragment midpoints to a continuous quantification along the genome (Boyle et al. 2008). To capture the benefits of regressing precisely the effects that we wanted (intercept and enrichment by IP), we adapted F-Seq's kernel density approach to place a kernel at the midpoint of each sliding window, then re-estimate density at these points corresponding to the observation windows. For this correction to the quantifications due to noise, we used a Gaussian kernel with sigma = 40 bp, specific to the observations with window size of 40 bp. These corrected responses are repeated to fill the 20 bp step size, aligned and averaged in the case where we are profiling multiple genes (as a line plot), and then the $L_2$FE is calculated using arithmetic on the intercept and IP responses.

Again, for zooming out to whole-chromosome view, we wanted our line plot of enrichment on the chromosome to average over many kb, because only trends of this bandwidth will be evident, and finer details will appear as uninterpretable noise. Thus, to interpret the $L_2$FE regression coefficient (500 bp window observations) at a zoomed-out view of more than 1 Mb, we applied a Gaussian kernel with sigma = 1 kb to the sequence of the regression effects filled by step size. The $L_2$FE after this step was plotted for the dm6 first 7 reference sequences (which cover the chromosomes except for some telomeric and centromeric heterochromatin). Although applying this bandwidth helped with the cleanliness and interpretability of genome-wide line plot graphics (where one pixel of the graphic steps more than 1 kb), we also checked the $L_2$FE values (which originally use a 500 bp-wide observation) for their distribution and found that they can be used directly for downstream analysis (arithmetic mean and *t*-test). Tests for the suitability of ChIC-ChIP $L_2$FE coefficients for downstream analysis include a violin plot (visually inspecting the distribution for normality), 2D binning (two cell types may show a bivariate Gaussian), and Q-Q plot of the residuals (regressing out each reference sequence and each pericentromere's $L_2$FE enrichment). An alternative analysis could use a Wilcoxon test, which we found to be similarly strong at rejecting the null hypothesis in many places but is much less interpretable (with *t*-statistic) than a *t*-test.

<u>Integrated $L_2$FE</u>

Published datasets used for the chromatin analysis include:

modENCODE Kc, H3K4me3: GSE45088; H3K27me3: GSE45083; H3K9me3: GSE27796

modENCODE S2, H3K4me3: GSE 20787; H3K27me3: GSE 20781; H3K9me3: GSE20794

ChIP-on-chip was first scored as an intensity value, and then the microarray spots (which corresponded to DNA strings taken from the dm3 reference) were scaled to have zero mean and unit variance *(Z*-scoring). The distribution of *Z*-score of Intensity was skewed (not normal), even counting the original microarray spots and before lifting and averaging sliding windows of the dm6 reference. We found that the ChIC-ChIP $L_2$FE coefficients of GSC-like and CySC-like cells were approximately normally distributed, with a standard deviation (at $\log_2$ scale) of approximately 1. Thus, to integrate ChIP-on-chip observations to have a similar distribution as ChIC-ChIP $L_2$FE, we simply needed to replace the ChIP-on-chip intensities with the rank of the intensity (at all sliding windows on the genome), and then look up the ranks (scaled to be between 0 and 1, exclusive) on the normal distribution quantiles. This yielded an approximately normal histone mark intensity score, at approximately the same scale as the GSC-like and CySC-like cells' $\log_2$ coefficients (Integrated $L_2$FE).

<u>Chromatin Classification</u>

As we will turn to the Kc167 cell line genomics (for some of our comparisons and for the quantification of trends in chromosome regions), we can apply a model fitted to the Kc167 epigenomics. Our goal is to classify the pericentromere in a manner that explains some of the between-region variance in a cell type of major interest to our experiments. The Kc167 hidden Markov model inference, where the emissions are the enrichment of 53 broadly selected chromatin components, labels domains that likely belong to a distinguishable enrichment profile of chromatin components (Filion et al. 2010). On each dm6 autosome reference sequence (chromosome arm), we labeled the run of at least 50 kb of classic (including pericentric) heterochromatin nearest the centromere, and extending towards the sparsely covered chromatin towards the centromere (which generally did not have a predicted chromatin class) as the pericentromere region. The Kc167 epigenomic euchromatin-pericentromere border is an adjustment relative to the pericentromere definition provided with the *D. melanogaster* genome and is highly like the embryo epigenomic euchromatin-pericentromere border in location. In particular, the coordinates of the pericentromere can be checked for gene silencing, as the gene *clamp* is on in our 5 clusters (scRNA-seq) and is not within our pericentric coordinates for the testis epigenome. Placing "on" genes just outside of the pericentromere is a finding credited to checking for evidence of pericentric chromatin marks in a relevant cell line.

Cell-Type-Specific Chromatin

We looked at the regions of the chromosome, and now we turn to marking regions of chromatin as enriched for a histone mark in either specifically GSC-like cells or specifically CySC-like cells. These chromatin regimes are summarized at the chromatin classification (arm & pericentromere) level as heatmaps. Cell-type-specific chromatin in the 11 chromosome regions plus rDNA is detected in a heatmap of GSC-like $L_2FE$ on the *x*-axis and CySC-like $L_2FE$ on the *y*-axis. The $L_2FE$ scores tested here are regressed on a rectangular sliding window of 500 bp width (several times the nucleosome spacing), making these tracks of $L_2FE$ score being summarized highly similar to the F-Seq-smoothed genomics plots. The cell-type-specific chromatin quantification is the proportion of chromatin that lies meaningfully inside the GSC-specific quadrant or the CySC-specific quadrant (where the median-normalized $L_2FE$ estimates have different signs and are not near zero). We chose a requirement that $|L_2FE| >= 0.2$ based on standard error estimates of $L_2FE$ being almost 0.2 across the ChIC-ChIP experiments. Thus, we are counting a proportion of the chromatin that possesses $L_2FE$ scores that are meaningfully distinct in the GSC-like and CySC-like cells. For this analysis, we produced GSC-CySC $L_2FE$ correlation heatmaps as well as a quadrant summary (2 quadrants) after filtering by this $|L_2FE|$ threshold.

The quadrant summary is tested against the null hypothesis of uniform GSC-Specific and CySC-Specific chromatin percentages (Supplemental Table 4). The test used is a permutation test, simulating assigning the labels at a 50/50 likelihood to the total number of chromatin windows that are either GSC-Specific or CySC-Specific. There is an exact probability distribution under the null hypothesis (Binomial distribution). This yields a p-value for the null hypothesis that the histone mark is not specifically enriched in either cell type in the chromosome region, which is then adjusted using the Benjamini-Hochberg procedure to determine statistical significance.

*Repli-seq QC*

Repli-seq sequencing results were aligned to the dm6 genome using Bowtie 2 (version 2.4.5) (Langmead and Salzberg 2012). For Repli-seq, in single-end technology, each read is taken to be a fragment of length 100 bp, as the step size (1 kb) is several times larger than the fragmented DNA and so the exact coordinate estimate is not necessary. No duplicate filtering is performed on single-end reads that have the same alignments. For paired-end technology, proper pairs that do have the same alignment are likely to be PCR duplicates, so markdup (followed by filtering out of fragments assigned the duplicate tag) is performed. Next, the midpoints of reads or fragments are counted in bins of size 1 kb, so the same fragment is not counted in adjacent bins (as in the case of sliding windows).

The success of the Repli-seq experiment rests on the fractions being enriched (or depleted) for distinct broad regions of nascent DNA. To assess every sample, the bins' midpoints and counts are treated as observations to be smoothed, either using LOESS or a Gaussian kernel with sigma = 50 kb (the F-Seq approach with extremely broad density estimation across the genome). The bins (in rows) are bound together for all samples, and the columns, and then the rows, are unit-scaled and centered *(Z*-scored). Next, a PCA reveals whether every sample has good information content, or whether some middle fractions do not have much difference at all to the other fractions or to the line connecting the early and late fractions (they are a mixture of the earliest and latest chromatin which those two most extreme fractions can characterize without this middle fraction).

Repli-seq Regression

We parameterized the nascent DNA at a location by the mode and concentration of fragments (appearing in fractions of sorted cells). The Beta distribution is ubiquitous in Bayesian models, as it has the following properties: The Beta-distributed random variable is on a closed interval (given as [0, 1]), the distribution can be symmetric or skewed, and a prior can be given so that the posterior distribution is either uniform or unimodal. To form the model design, the Beta distribution with parameters $(\alpha, \beta)$ (two degrees of freedom) is cut according to the quantiles of the cells' DNA content quantifications (quartiles of the Beta function). For model likelihood, like Negative Binomial (Gamma-Poisson) regression, we take the fragment counts to follow a distribution of overdispersion counts (compound distribution). Like Negative Binomial regression, the scale parameter from the model is multiplied by a shape parameter $\theta$ (overdispersion). Like the GLM Gamma-Poisson implementation in R, we don't calculate the posterior distribution of $\theta$ , but we only make a maximum-likelihood estimate of this nuisance parameter and apply it to the model. Regression with Dirichlet-Multinomial likelihood differs from Gamma-Poisson regression in that the responses are structured (vector-valued), and overdispersion models the same total number of trials (fragments) but which could be shuffled into different outcomes (the cell fractions in the same biological replicate are not statistically independent).

Like Gamma-Poisson regression of high-throughput sequencing data, there is a random effect of sequencing depth in every HTS sample which we cannot control, and which is estimated for every HTS sample and at a genome-wide scale (FPKM normalization factor: one million divided by the product of the total count of fragments and the observation window's size in kb). The model responses for the fractions of cells, parameterized by $(\alpha, \beta)$, are multiplied by the FPKM normalization factors, and normalized to again sum to 1. If the samples in the regression design do not cover all percentiles (e.g. after QC, observing only the first quartile and last quartile for one replicate), then, once again, the vector with fewer entries is normalized to sum to 1.

The DNA replication pseudotime *t* is a parameter distributed with Logistic prior (Logistic Regression). The posterior expected value of *t* at a genomic location is our Repli-seq timing estimate for the cell type/condition. A nuisance parameter $\kappa$ explains the uniformity (lack of enrichment) or concentration (enrichment) of the nascent DNA in one or more fractions. The parameter $\kappa$ appears in the mode-concentration $(\sigma(t), \kappa)$ parameterization of the Beta distribution, which defines our nascent DNA fraction abundance. Predictions of the abundance of each nascent DNA fraction are passed to the likelihood function. Unlike Negative Binomial regression of HTS data, $\sigma(t)$ (the time scale) is in a fixed interval, and the prior for $\kappa$ can also be a uniform or truncated distribution, motivating us to carry out the Bayesian inference by integrating the prior times the likelihood, rather than by an approximation (as applied previously for quantifying DEGs). We chose a truncated normal prior for $\kappa$, on the interval [0.1, 10], with a mean of 2.5 and a scale of 1. To decide on a wide and effective truncated range of $\kappa$ for Repli-seq model fitting, we simulated possible Repli-seq fraction predictions with our parameters and with the R function pbeta. The parameter $\kappa$ could best be described as on a similar scale to the L2FE from the least abundant to most abundant nascent DNA fraction being predicted. Thus, we chose a range of model parameters wider than what we empirically observed in the 4 Repli-seq experiments studied (ranging from nearly uniform HTS abundance between fractions corresponding to small $\kappa$, to testing the likelihood of an L2FE between at least some of the fractions of 10 or more).

Bayesian inference proceeds as follows. The sigmoid function $\sigma(t)$, as applied in Logistic Regression, takes *t* from the domain of the real number line to the range $(0,1)$. The Logistic Regression produces model responses (percentages) according to the Beta distribution parameters: $(\alpha, \beta) = \left(1 + \kappa\sigma(t),\ 1 + \kappa\left(1 - \sigma(t)\right)\right)$. The prior times the likelihood will be integrated in polar coordinates in terms of $(\alpha, \beta)$. The integral is evaluated at points on a polar grid of $1.2°$ by 0.1. The nuisance parameter $\kappa$ can be integrated out immediately, as the posterior distribution $P(t)$ is the track (computed on sliding windows) that we analyze.

Finally, we reported the Replication Score using a tanh-like layer, and negated: $1 - 2\sigma(E[t])$. This stylistic choice highlights early-replicating, potentially accessible, chromatin with a positive score, while making late-replicating chromatin negative.

The regression design was realized using more individuals than the two biological replicates per cell type. We reviewed the Repliseq pileups summarized as bins of size 1 kb (one fragment will generally appear in exactly one bin), testing the autocorrelation at a lag of 1 kb. The autocorrelation $R^2$ was between 0.8 and 0.9: the DNA replication regime does not actually change at this distance, but adjacent bins can be distinctly informative. The 3 adjacent bins (observations totaling 3 kb for Repli-seq) were each quantified separately and treated as independent observations (2 replicates and 6 vector-valued observations for each cell type). The greater number of observations to regress will aid in explaining (relying on the nuisance parameters in the model) how the low input of the Repliseq experiment influences the signal-to-noise ratio.

Nested Differential Replication Peak Calling

We introduce Bayesian hypothesis testing, and nested peak calling, to our Repli-seq regression method. Consider a null hypothesis, for Repli-seq logistic coefficients and, that these parameters are held to have the same posterior distribution ($P(t_1 = t) = P(t_2 = t)$), and an alternate hypothesis where the two coefficients (cell type replication scores) have different posterior distributions. The posterior distributions $P(t_1)$, $P(t_2)$, have a domain over the real number line, and the integral: $\int P(t_1 = t) \cdot P(t_2 = t)dt$ is an inner product for this space of posterior distributions. The integral produces an odds ratio, between 0 and 1, of the null hypothesis to the alternate hypothesis. The odds ratio will be reported as the reciprocal of the integral and is a Bayes Factor (odds ratio for the differential replication hypothesis of interest). The null hypothesis is rejected for a Bayes Factor larger than a threshold, which we set at Bayes Factor at least 100 and which we call a Bayesian two-star significance level. The bins of Repli-seq (1 kb step) are each colored by whether the null hypothesis is rejected for a pair of coefficients (cell types). Next, color is filled in gaps (rejecting the null hypothesis for regions) where the Bayes Factor was locally less than 100 but is a gap of length at most 10 kb, flanked by bins with Bayes Factor at least 100. Finally, contiguous color (a nested peak) must have a width at least 20 kb. The nested peak contains bins where the model fit and the difference in replication coefficients could vary substantially. We classify the nested peak by the sign of the arithmetic mean of the most negative replication coefficient difference and the most positive replication coefficient difference (which assumes that the bins in the nested peak predominantly have one sign of difference between cell types).

Nested peaks' widths, for one direction of between-cell-type difference, are summed in each chromosomal region. These sums were divided by a "replication regime width" (50 kb) followed by a ceiling function, to count individual observations where we rejected the null hypothesis for a regime, without chopping the genome overly finely. Then, Fisher's exact test is applied to this continuous-to-discrete contingency table, to explore whether our chromosomal regions do help us to characterize cell-type differences in replication choreography.

Repli-seq Cell Type Comparison

The datasets used in our Repli-seq Cell Type Comparison are as follows:

Kc167, GSE41349

S2, GSE41350

modENCODE Orc2 ChIP meta-peaks: GSE27123

For between-Repli-seq plotting and Euclidean distance specifically, we introduced LOESS local regression of the timing scores along each reference sequence. The LOESS parameter applied was span = 0.025. LOESS removes the high-frequency variance at small steps on the genome and has already been needed to interpret Repliseq scoring (Lubelsky et al. 2014). The application of LOESS happens after we demonstrate that we can score replication timing (RT) at a fine level on the bins of low-input fragments (only looking at 3 kb of the genome) and now need to smooth the variance that we observed at this fine level to view peaks and valleys in RT genome-wide. LOESS is a necessary step before genome-wide Pearson correlation, so that the total variance in the RT is not excessive and dynamic between experiments, but the variance represents plausibly different replication regimes at a reasonable distance (smoothing tens of kb). The LOESS of the RT predictions can be applied immediately to produce a Euclidean distance matrix between cell types, yielding a highly interpretable hierarchical clustering of the cell types. The LOESS plot is also paired up with the nested differential replication peak calling in a graphic. Although they are different secondary analyses (the LOESS is discrete to continuous, whereas the nested peaks look within the original step size and only connect within RT peaks at a fine level), both analyses highlight the peaks and valleys of the RT regime.

Lining up cell types in order by their hierarchical clustering, and summarizing the original RT estimates, can again confirm whether the definitions of the pericentromeres are appropriate for the experiments. To quantitatively compare the cell types, we did not plot the Euclidean distance matrix, but instead *Z*-scored the variance between the cell types at each genomic location. Mean *Z*-scores reveal how the cell types hierarchically differ at the pericentromeres, chromosome arms, and other loci, and clearly show when the full extent of some of the chromosome arms are overall statically replicating between the cell types. For downstream analysis, we centered the *Z*-scores of each cell type to have zero mean, and then weighted (multiplied) their contribution to the $R^2$ (Pearson) by the reciprocal of the length of the assigned region (11 arms and pericentromeres, and a single unit length rDNA locus) that they belong to (the pericentromeres are shorter sequences and aid in characterizing cell-type differences). For simplicity, we tested the Pearson correlations of the vector of the 12 mean *Z*-scores, and as we found that this produced the same signs of the Pearson correlations, we used this summary matrix for correlation analysis (Correlation of *Z*-Scores). This fully characterizes cell-type differences in terms of chromosome-scale shifts in replication priority, and complements the nested peak calling, which proceeded from the unit step size upwards.

Statistics and Reproducibility

The regression coefficients (ChIC-ChIP $L_2$FE, or RT value logit) were approximately normal (method used was visually inspecting a violin plot with a small bandwidth parameter) and were subjected to a Welch's *t*-test (`t.test`). GSC-like and CySC-like observations are only paired (`paired = TRUE`, disabling the Welch's option) for the regression coefficient on all FlyBase transposable elements. For every panel or table, the p-values were adjusted by Holm's method to control the family-wise error rate (`p.adjust`). Cell-type specific chromatin is tested using a permutation test assuming a 50/50 Bernoulli distribution (`pbinom`), and the differential replication program (summarized by the replication regime width) is split into two columns (according to the sign of the cell-type effect) and tested with Fisher's exact test (`fisher.test`). Significant differences are marked at four significance levels (* $P < 0.05$, ** $P < 0.01$, *** $P < 0.001$, **** $P < 0.0001$).

For scRNA-seq and for Repli-seq, a posterior distribution is inspected directly (from the heavy-tailed prior model for scRNA-seq, and from polar integration for Repli-seq) for Bayesian hypothesis testing. The scRNA-seq local false sign rate has a correspondence to Wald's test (checking the cumulative distribution function at 0 to yield the p-value), but the Bayesian treatment of the heavy-tailed L2FC parameter as a random effect already controls for multiple hypothesis testing, so a method-specific FDR-controlling value (an *S*-value) is produced. As many *S*-values are less than 0.05 (we have more than enough observations of single cells to reject the null hypothesis), only one significance level is tested for scRNA-seq: *S*-value < *0.0001*. For Repli-seq, in the joint distribution of both cell types' timing, we integrate only the part of the distribution where the cell-type logistic coefficient is 0, producing an Odds Ratio. Like the scRNA-seq hypothesis testing, only one significance level is tested for coloring the genome by replication program, and it is also stronger than the one-star level: Odds Ratio >= 100.
